# A glimpse of the paleome in endolithic microbial communities

**DOI:** 10.1101/2022.09.23.509128

**Authors:** Carl-Eric Wegner, Raphaela Stahl, Irina Velsko, Alex Hübner, Zandra Fagernäs, Christina Warinner, Robert Lehmann, Thomas Ritschel, Kai U. Totsche, Kirsten Küsel

**Author notes:** Corresponding author: Kirsten Küsel.

## Abstract

The terrestrial subsurface houses a significant proportion of the Earth’s microbial biomass. Our understanding about terrestrial subsurface microbiomes is almost exclusively derived from groundwater and porous sediments. To obtain more insights about endolithic microbiomes and their metabolic status, we investigated rock samples from the vadose zone, fractured aquifers, and deep aquitards. Using methods from paleogenomics, we recovered sufficient DNA for metagenomics from rock specimens independent of porosity, lithology, and depth. We estimated between 2.81 and 4.25 × 10^5^ cells × g^−1^ rock. DNA damage patterns revealed paleome signatures (genetic records of past microbial communities) for three rock specimens from the vadose zone. The taxonomy and functional potential of paleome communities revealed increased abundances of chemolithoautotrophs, and a broader metabolic potential for aromatic hydrocarbon breakdown. Our study suggests that limestones represent ideal archives for genetic records of past microbial communities, due to their specific conditions facilitating long-term DNA preservation.

## Introduction

The subsurface harbors a significant portion of the Earth’s microbial biomass and contributes to global biogeochemical cycling [1–3]. The difficulty of access impairs estimating global subsurface biomass, activity, and biodiversity, especially in the continental biosphere. A comprehensive compilation of cell count measurements suggested that there are approximately 2 to 6 × 10^29^ cells in the continental subsurface [3]. Biomass estimates are exposed to significant uncertainties due to poorly understood parameters such as the ratio of surface-attached to pelagic groundwater cells, for which assumptions range between 1 and 10000. total organic carbon content and groundwater cellular abundances have been shown to be poor predictors for biomass and biodiversity [1–4].

Groundwater and other aqueous sample material are keys [4–8] for studying subsurface microbiomes, but only provide limited information regarding surface-attached or endolithic microbes inhabiting rock matrix pores. Microbial communities inhabiting the subsurface have been studied predominantly in porous, unconsolidated sediments, for example alluvial aquifer systems [9–12]. The bedrock itself has been rarely investigated [13–15]. Similarly, the vadose zone, the shallow unsaturated bedrock zone more connected to surface habitats [16, 17], has received little attention. Water saturation and nutrient supply, both controlled by relief position, rock properties (i.e. porosity, permeability, fracture network, composition), and groundwater quality and circulation patterns control subsurface microbial life [16, 18]. The subsurface endolithic microbiome consists of subsurface specialists that prefer particular lithologies [19, 20], long-term descendants of organisms that colonized sediments during deposition [21], surface immigrants transported by fluid flow over geological time [21, 22], and invaders introduced as a result of human activities, such as drilling or flooding. Continental subsurface habitats are viewed as energy-starved systems. They suffer from a lack of electron donors, electron acceptors, carbon, and nutrients [19], and are characterized by generation times in the range of thousands of years [23]. Ancient sedimentary carbon might represent a significant source of carbon for microorganisms in subsurface rock environments [24–26]. Part of these carbon compounds can be still metabolized, diffuse from aquitards into aquifers [27] and from less permeable into more permeable layers where they drive microbial activity [9, 11].

Because of the low amounts of microbial biomass and the challenge of recovering it, 16S rRNA gene amplicon studies from rock core material have been the primary means of investigating microbial community composition. These surveys provide limited insights into the metabolic potential of organisms, and by default do not allow discrimination between living, potentially active, and dead cell material. Advances in paleomicrobiology, achieved through distinguishing “ancient” and “modern” DNA by high-throughput sequencing and DNA damage pattern analysis, are potential door openers for subsurface microbiology. Similar to hard tissue samples (bone, teeth, shells) [28–31], carbonate rocks contain calcium carbonates and calcium phosphates, which could adsorb or encase extracellular, “ancient” DNA by neutralizing negative charges present in the DNA backbone and the mineral surface by bivalent calcium cations [32]. We hypothesized that limestone/marlstone parent material would allow the recovery of metagenomic DNA (mgDNA), which could be sufficient to gain insights into the genomic potential of endolithic microbes.

In this study, we adapted methods from microbial archaeology and paleogenomics for mgDNA recovery, used comprehensive wet- and dry-lab control measures to minimize the risk of contamination, applied metagenomics, and analyzed DNA damage patterns. The goal was to assess endolithic microbial biomass and use DNA damage as a proxy to distinguish DNA from intact and potentially alive cells from the paleome, the genetic remains of past microbial communities [30]. In addition, we aimed for decoding the taxonomic compositions and metabolic potentials of the endolithic microbiomes to understand how these communities are or were adapted to a life in consolidated rocks.

## Materials and Methods

### Bedrock sampling and sample preparation

We collected fractured bedrock from Upper Muschelkalk marine deposits (Germanic Triassic) in the groundwater recharge area of the Hainich low mountain range, as well as from isolated equivalents in the center of the Thuringian Syncline (both central Germany). Sampling was done during the construction of groundwater monitoring wells (Hainich CZE: 2011, 2014; samples: H13-17, H22-8, H22-30, KS36-H32, CM1-H32) and during the INFLUINS exploratory drilling (EF1/2012: 2013; samples: INF-MB2, INF-MB3). Measures to minimize contaminations included utilization of washed, de-rusted, steam-cleaned drill pipes, as well as ethanol-washed PVC liners in the Hainich CZE. Selected core segments of drill cores, recovered with rotary drill rigs (mud-rotary wireline), were immediately wrapped in sterile plastic bags, and transported on dry ice until storage in deep freezers (−80°C). Subsamples of bedrock matrices for DNA-extraction were prepared by fast hydraulic splitting of still frozen drill cores, under removal of the outer parts of the core segments. Subsamples for X-ray micro-computed tomography analysis (13 mm plugs, vertical orientation) were prepared with a drill press. Samples for carbon analysis were extracted from directly adjacent rock and also used for rock typing.

### Rock typing/characterization, pore classification and analysis of carbon fractions

The rocks were classified based on stereoscope inspection, carbonate test (HCl 10%), and analytical carbon measurements by applying the Dunham [33] and a mudrock classification scheme [34]. Porosity types and pore sizes were classified as described previously [35, 36]. Milieu indicators, including weathering colors and secondary pore mineralizations (Munsell colors), and derived oxicity rating were determined by stereoscope inspection, and also contrasted against characteristics of the core segment and borehole/well. The contents of total carbon and organic carbon (TOC) of the rock samples were determined on homogenized duplicate subsamples (~1.6 mg) of ~30 g ground rock using an elemental analyzer (Euro EA, HEKAtech, Wegberg, Germany). The OC was calculated as the difference from total carbon measurements released under combustion at 950°C and 600°C.

### X-ray micro-computed tomography (X-ray μCT)

The three-dimensional structure of the plugs was assessed non-destructively by X-ray μCT (Xradia 620 Versa, Zeiss, Jena, Germany). Each plug was scanned in 1601 projections to give a full 360° rotation at 0.4× magnification and an exposure time of two seconds per step using X-rays produced with 80 kV and 126 μA. Tomographic reconstruction yielded a three-dimensional grayscale image with 1024^3^ voxels at a resolution of 25.99 μm with automated removal of ring artifacts and beam hardening. Images were cropped to remove boundaries, denoised with non-local means filtering [37] and binarized into pore space and solid by manual thresholding using Fiji (ImageJ v. 1.51) [38]. The pore sizes were calculated from binarized images using the maximum inscribed sphere algorithm implemented in the BoneJ plugin [39] in Fiji. The volumetric pore size distribution was derived from the histogram of resulting images, while total X-ray μCT visible porosity was derived directly from the histogram of binarized images. Connected pore space of binarized images was assessed assuming 26-connectivity and visualized by randomly assigning a color to each set of voxels belonging to the same region.

### Protocols for DNA extraction and sequencing library preparation

We adapted protocols routinely used for ancient DNA preparation for downstream sequencing, which are all available from protocols.io (https://dx.doi.org/10.17504/protocols.io.bvt9n6r6). We reference the respective protocols in the following sections and describe them for the sake of completeness. The bench protocols available on protocols.io include detailed lists with respect to needed equipment and reagents, as well as necessary precautions.

### DNA extraction

DNA extraction from rock samples was performed by modifying a protocol originally designed for recovering ancient DNA from dental calculus (https://dx.doi.org/10.17504/protocols.io.bidyka7w). Metagenomic DNA was extracted from either 2.5 g of rock powder obtained using a dental drill or 2.5 g of rock pieces obtained by chipping rock material. To decalcify the samples, the rock material was rotated in EDTA (0.5 M, pH 8.0) for up to 10 days (rock pieces, rock powder 5 days) at 37 °C before being concentrated down to a volume of 1 mL using Amicon ® ultra centrifugal filtering units (MWCO 30 kDa and 10 kDa). Concentrated samples were mixed with 1 mL of extraction buffer (EDTA pH 8.0, 0.45 M; Proteinase K 0.025 mg/mL) and rotated overnight at 37°C. Samples were spun down and subsequently mixed with 10 mL of binding buffer (guanidine hydrochloride, 4.77 M; isopropanol, 40% [v/v]) and 400 μL sodium acetate (3 M, pH 5.2). Samples were transferred to a high pure extender assembly from the High Pure Viral Nucleic Acid Large Volume kit (Roche, Mannheim, Germany) and centrifuged for 8 min with 1,500 rpm at room temperature. The column from the high pure extender assembly was removed, placed in a new collection tube and dried by being centrifuged for 2 min with 14,000 rpm at room temperature. 450 μL of wash buffer (High Pure Viral Nucleic Acid Large Volume kit) were added and samples were centrifuged for 1 min at 8,000 × g at room temperature. This washing step was repeated once and columns were dried afterwards by centrifugation. DNA was eluted into a siliconized tube by adding 50 μL of TET (0.04% Tween 20 in 1 × Tris-EDTA [pH 8.0]), incubating samples for 3 min at room temperature, and centrifugation for 1 min 14,000 rpm at room temperature. The elution step was repeated once and the pooled eluate was stored at −20 °C until further processed. All outlined steps were carried out in the ancient DNA lab of Max Planck Institute for the Science of Human History (MPI-SHH) to reduce the risk of contamination with modern environmental DNA. Blank extractions were carried out alongside the sample extractions, using identical steps, with the exception that water instead of rock material was used as input material. DNA concentrations were determined using a Qubit® fluorometer and the DNA high-sensitivity assay (ThermoFisher, Schwerte, Germany). Cell number estimates were calculated by dividing the amount of extracted DNA per gram rock material by the approximate mass of one prokaryotic genome, assuming a molecular weight per base pair of 618 Da (g/mol) [40] and a genome length of 3 Mbp.

### Library preparation

Anticipating that extracted metagenomic DNA could contain both severely fragmented ancient DNA and high molecular weight modern DNA, we first used a Covaris M220 ultrasonicator to shear any high molecular weight DNA present to a maximum length of 500 bp prior to library construction. This ensured that all DNA present in the DNA extract would be suitable for library construction. We then used a library construction protocol (https://dx.doi.org/10.17504/protocols.io.bakricv6) that is specifically designed to be compatible with degraded and ultrashort DNA fragments [41]. Metagenomic DNA samples were blunt end repaired by mixing 10 μL of DNA with 40 μL of a mastermix containing NEB buffer no. 2 (1×), ATP 1 mM, BSA 0.8 mg/mL, dNTPs 0.1 mM, T4 PNK 0.4 U, and T4 Polymerase 0.024 U. Samples were incubated for 20 min at 25 °C, followed by a 10 min incubation step at 12 °C. Blunt end repaired samples were subsequently purified using the MinElute Reaction Clean-up Kit (Qiagen, Hilden, Germany). Samples were finally eluted in 20 μL of the elution buffer containing 0.05% Tween20. 18 μL of eluted samples were mixed with 21 μL of a mastermix containing Quick Ligase buffer (final concentration 1 ×) and a mix of adapters (0.25 μM). Next, 1 μL of Quick Ligase (5 U) was added and libraries were incubated at 22 °C for 20 min. Reactions were again purified using the MinElute Reaction Clean-up Kit. Samples were eluted using 22 μL elution buffer. The adapter fill-in reaction was performed in a final volume of 40 μL. The reaction mix consisted of a 20 μL eluate and a 20 μL mastermix containing isothermal buffer (final concentration 1 ×), dNTPs (0.125 mM each), and Bst polymerase (0.4 U). Reactions were incubated for 30 min at 37 °C, before being incubated at 80 °C for additional 10 min to inactivate the polymerase. Before being further processed, libraries were quality-checked by quantitative PCR (qPCR). Dilutions of the libraries (1:10) were mixed (1 μL template), with 19 μL of a mixture containing DyNAmo mastermix (final concentration 1 ×) and IS7 and IS8 primers (1 μM). The thermal profile was 10 min at 95°C, 40 cycles of 30 s at 95°C, 1 min at 60°C, 30 s at 72°C, followed by a melting curve (60-95°C). Libraries were subsequently indexed (https://dx.doi.org/10.17504/protocols.io.bvt8n6rw) and amplified (https://dx.doi.org/10.17504/protocols.io.beqkjduw) as outlined in the referenced protocols. Libraries were equimolarly pooled and sequenced on an Illumina NextSeq 500 instrument in paired-end mode (2 × 150 bp) using v. 2.5 chemistry. The sequencing depth ranged between 2.24 and 4.81 Gbp (**Table S1**). All outlined steps were carried out in the ancient and modern DNA clean rooms of the MPI-SHH to reduce the possibility of contamination. Library blanks were prepared alongside the sample extractions, using identical steps, with the exception that water instead of rock material eluate was used as input material.

### Sequence data pre-processing

Quality parameters of raw sequencing data were assessed using *FastQC* (v. 0.11.8) [42]. Adapter and quality trimming was done with *bbduk* (v. 38.22) [43] (settings: qtrim=rl trimq=20 ktrim=r k=25 mink=11) using its included set of common sequence contaminants and adapters. Trimmed sequences were subsequently subjected to taxonomic profiling and metagenome assembly and binning.

### Taxonomic profiling

Trimmed sequences were taxonomically profiled using *kaiju* (v. 1.7.3) [44] and *diamond* (v. 2.0.7.145) [45, 46]. *Diamond* was used for the taxonomic assignment of trimmed, and paired-end assembled (with *vsearch* (v2.14.1) [47]) sequences, while *kaiju* was used for the taxonomic assignment of assembled contigs. For *kaiju*, sequences were translated into open reading frames, which were used for string matching with the implemented backward-search algorithm based on the one that is part of the Burrows-Wheeler transform [48, 49]. *Kaiju* was run in greedy mode with up to 5 allowed mismatches (-a greedy -e 5). *Diamond* searches were done in sensitive mode applying an E-value threshold of 0.0001 (-e 0.00001 -c 1 --sensitive). Database hits were annotated making use of the LCA algorithm implemented in *megan* (v. 6.21.1) [50, 51] with default settings. NCBI nr [52] was used as the reference database for taxonomic profiling (*kaiju,* nr_euk release 2020-05; *diamond,* custom built database based on NCBI nr retrieved from NCBI in 2020-03). Taxa representing contaminants on different taxonomic levels were identified using taxonomic profiles obtained from *diamond* and *decontam* (v. 1.1.1) [53] based on prevalence and frequency in true samples and extraction and library blanks.

### Metagenome assembly and binning

Metagenome coverage was estimated based on k-mer redundancy using *nonpareil* (v. 3.303) (-T kmer) [54, 55]. Trimmed sequences were assembled into contigs with *megahit* (v. 1.2.9) (default settings) [56] and *metaSPADES* (v. 3.13.0) (--only-assembler) [57, 58]. Due to better performance we used the megahit assemblies for all subsequent steps. Contigs longer than 1 kb were kept and quality-controlled sequences were mapped onto these contigs using *bowtie2* (v. 2.3.4.1) [59] (--no-unal). Resulting .sam files were converted into .bam files and indexed with *samtools* (v. 1.7) [60]. Contigs and indexed mapping files were used for manual metagenomic binning using *anvio* (v. 6.2) [61] based on sequence composition and differential abundance. The completeness, redundancy, and heterogeneity of bins was assessed with *checkm* (v. 1.1.2) [62]. Bins were taxonomically assigned using *gtdb-tk* (v. 0.3.2) [63].

### Functional annotation

Functional profiling of trimmed sequences was done with *humann* (v. 3.0) [64] using precompiled Uniref50 and Uniref90 protein databases (release 2019-01) and applying default settings. The resulting gene families table was regrouped to KEGG orthologies, normalized to copies per million (CoPM), and summarized with respect to pathways and functions of interest.

### DNA damage pattern analysis

Using assembled contigs and the output from mapping trimmed sequences back onto the contigs, DNA damage patterns were identified and analysed using *mapdamage* (v. 2.2.1) [65, 66] and *pydamage* (v. 0.50alpha) [67]. The output from *mapdamage* was ultimately used as it provides metrics with respect to all possible DNA damage-related substitutions. DNA damage pattern analysis was also done for selected subsets of the assembled contigs based on taxonomy (assigned with *kaiju*).

### Figure generation

Figures were prepared using the R packages *ggplot2* (part of *tidyverse*) (v. 1.3.1) [71] and *ggpubr* (v. 0.4.0) (https://rpkgs.datanovia.com/ggpubr/index.html) and finalized with *inkscape* (https://inkscape.org/).

### Data availability

Sequence data were deposited at the European Nucleotide Archive under BioProject number PRJEB52959.

## Results

### General sample characteristics and porosity analysis

We analyzed five bedrock samples from the vadose zone of a low-mountain range groundwater recharge area (Hainich Critical Zone Exploratory (CZE)) from depths between 9-33 meters below ground level (mbgl), and two samples from deep isolated aquitards with similar stratigraphic position and lithology (INFLUINS deep drilling) from depths 285 and 296 mbgl. The rock samples, representing the thin-bedded marine alternations of mixed carbonate-siliciclastic rock that form widely-distributed fractured-rock aquifers, range from argillaceous marlstones to bioclastic limestones with a broad range of porosity (**Table 1**). Three samples showed pores bigger than 0.02 mm (**Figure S1**) with volumetric fractions of 0.9% (INF-MB3), 2.4% (H22-30), or 8.9% (H13-17). INF-MB3 (**Figure S2**) showed a distribution of pores within 0.02-0.28 mm, which appeared at homogeneously distributed, but disconnected locations. The pore space in H22-30 (**Figure S3**) also shows several disconnected pores, but includes fractures and carbonate dissolution features that span large parts of the entire sample. The pore size distribution is slightly higher in the range of 0.02-0.52 mm. With pores in the size of 0.02-1.58 mm, H13-17 featured large macropores (**Figure 1**) from intensive carbonate dissolution that connect most of the internal pore space. H22-8 consisted of dense rock and showed only a single fracture in a size near the μCT limit of detection (**Figure 1**) that impeded a meaningful quantification. The other three rock samples did not show any pores above 0.02 mm, reflecting very dense rock matrices (**Figures S4-S6**). The macroscopic inspection revealed the presence of secondary Fe-minerals in large dissolution pores in two limestone specimens, also representing connected matrix habitats in the main aquifer (Trochitenkalk formation) (**Figure 1A + E**, **Table 1**). The total carbon content ranged between 5.53 ± 0.18 (H22-8) and 12.39 ± 0.17% (H32-KS36). The organic carbon content was, with the exception of CM1-H32 (8.17 ± 1.49%), below 3%.

**Table 1.** Origin and contextual data with respect to processed rock samples. Pandora DB refers to the internal sample database of the MPI SHH/MPI EVA. 1 = Dunham limestone classification after Wright (1992) [33] and mudrock classification (after Hennissen et al. 2017 [34], modified), 2 = (Visible) Carbonate porosity class after Ahr et al. (2005) [35]: apparent genetic factors: S (depositional; i: interparticle), D (diagenetic; d: dissolution; p: replacement; r: reduced; e: enhanced), F (fracture), 3 = Pore size classes after Choquette and Pray (1970) [112]: mc (micropores, <1/16 mm, macroscopically invisible), sms (small mesopores, 1/16-1/2 mm), lms (large mesopores, 1/2-4 mm), smg (small megapores, 4-32 mm), 4 = based on μCT analysis, pwd = refers to rock powder samples, pc = refers to rock pieces samples.

| Sample ID | Pandora DB ID | Sampling depth (mbst) | Aquifer type | Water saturation (in situ) | Rock type<sup>1</sup>, genetic porosity<sup>2</sup>, pore size classes<sup>3</sup> | Milieu indicators (plug, Munsell colors) | Oxicity | Porosity (%) | Total carbon (%) | Organic carbon (%) | Estimated cell concentration (g<sup>-1</sup>) |
| --- | --- | --- | --- | --- | --- | --- | --- | --- | --- | --- | --- |
| H13-17 | SET004.B01<br>03 (pwd)<br>SET004.B02<br>03 (pc) | 14.34-14.48 | fracture/<br>karst | saturated | Limestone (packstone); D (e); mc, sms-lms | matrix: 2.5Y 6/2; secondary Fe-minerals in pores | oxic | 8.9 | 12.00±0.02 | 2.95 ± 0.69 | 3.62E+05 |
| H22-8 | SET005.A01<br>03 (pwd)<br>SET005.A02<br>03 (pc) | 13.61-13.70 | fracture | unsaturated | A: Delithified argillaceous marlstone; F, D (e); mc<br>B: Delithified calcareous mudstone; F, D (e); mc | A, matrix: 2.5Y 7/4; B, matrix: 2.5Y 7/2 | oxic to suboxic | n.a. | A: 5.52 ± 0.18<br>B: 8.60 ± 0.10 | A: 1.42 ± 0.47<br>B: 2.12 ± 0.26 | 4.25E+05 |
| H22-30 | SET004.A01<br>03 (pwd)<br>SET004.A02<br>03 (pc) | 32.55-32.80 | fracture | unsaturated | Limestone (oolithic packstone); D (e), F; mc, sms-lms | matrix: 2.5Y 6/2; secondary Fe-minerals in pores | oxic | 2.4 | 11.92 ± 0.08 | 0.73 ± 0.09 | 2.38E+05 |
| KS36-H32 | SET001.B01<br>04 (pwd)<br>SET001.B02<br>03 (pc) | 8.92-9.06 | fracture | unsaturated | Limestone (packstone); D (e); mc, sms-lms | matrix: 5Y 5/1; secondary Fe-minerals in pores | oxic | n.a. | 12.39 ± 0.17 | 0.59 ± 0.14 | 3.52E+05 |
| CM1-H32 | SET003.A01<br>03 (pwd)<br>SET003.A02<br>03 (pc) | 21.9-22.0 | fracture | saturated | Calcareous mudstone to limestone (wackestone); D (r); mc | matrix: 10Y 3.5/1 | oxygen-deficient | n.a. | 10.74 ± 0.05 | 8.17 ± 1.49 | 4.12E+05 |
| INF-MB2 | SET001.A01<br>03 (pwd)<br>SET001.A02<br>03 (pc) | 285.44-285.62 | aquitard | saturated | Calcareous mudstone; D (r); mc | matrix: 5GY 2.5/1 | anoxic | n.a | 8.61 ± 0.09 | 2.93 ± 0.07 | 2.94E+05 |
| INF-MB3 | SET002.A01<br>03 (pwd)<br>SET002.A02<br>03 (pc) | 295.71-295.87 | aquitard | saturated | Limestone (packstone to grainstone); D (r); mc | matrix: 5Y 6/1; bioclasts: N3 | anoxic | 0.9 | 12.28±0.01 | 0.86±0.01 | 2.94E+05 |

**Figure 1.**
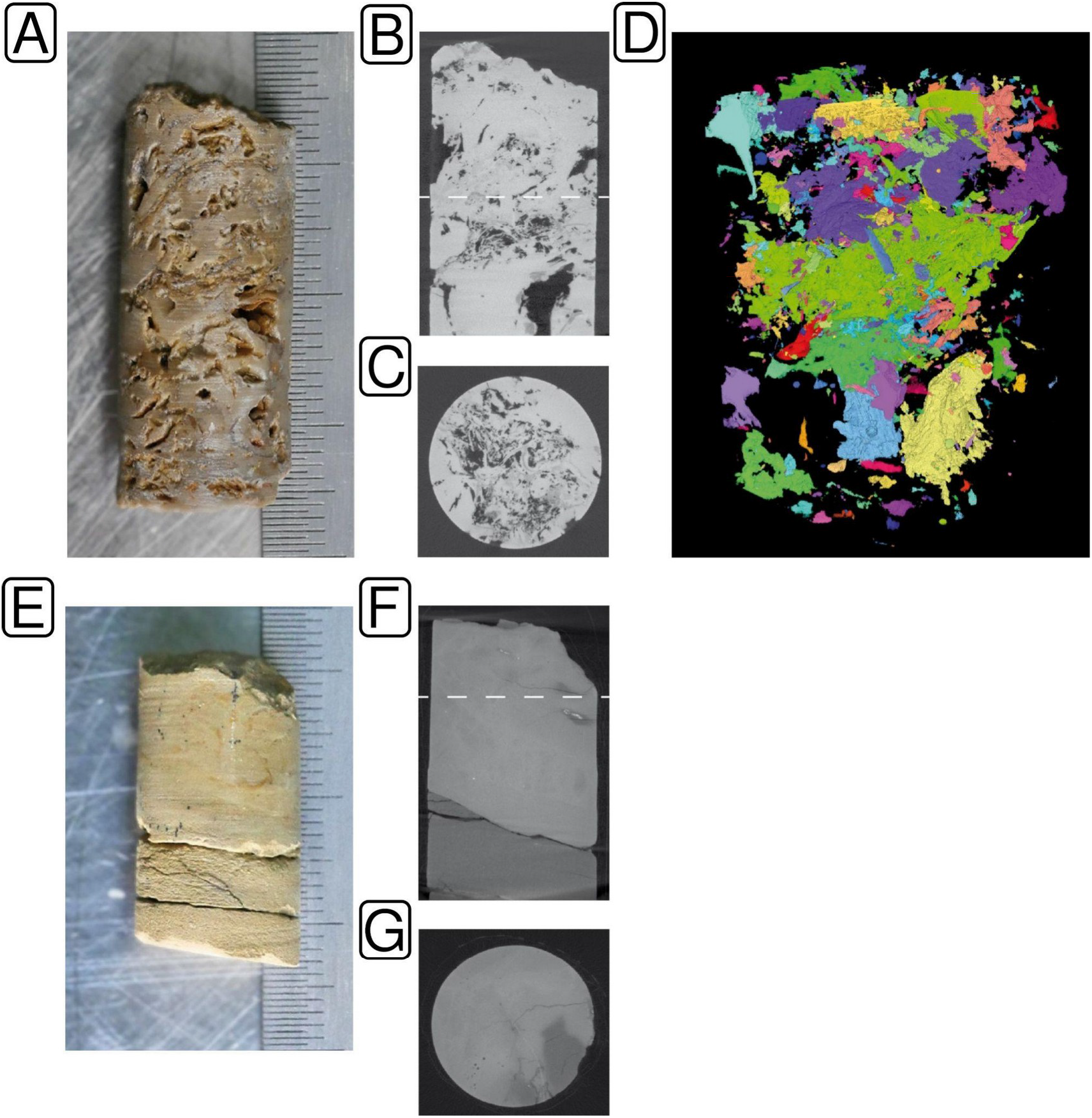
Pore space characteristics of samples H13-17 (A-D) and H22-8 (E-G) by μCT analysis. Moldic pores (up to large mesopores) dominate the packstone. Scale: 0.5 mm. Plug diameter 13 mm (A). Vertical section shows considerable porosity. The dashed line marks the position of C (B). Horizontal section (C). Reconstructed pore space. Colors mark parts of the pore system that are each connected by throats >26 μm (D). The plug comprises delithified siliceous marlstone (lower part) and delithified calcareous mudstone (upper part). Scale: 0.5 mm. Plug diameter 13 mm (E). Vertical section shows thin fractures (micropores). The dashed line marks the position of G (F). Horizontal section showing fine fractures and rare micro- to small mesopores. The matrix exhibits no pores connected by throats >26 μm (G).

### Recovery of metagenomic DNA (mgDNA) independent from the specimen

We were able to extract mgDNA from all rock specimens. DNA extractions yielded higher amounts from rock pieces than from ground rock powder (**Figure 2A**) with concentrations ranging between 0.011 and 0.051 ng × μL^−1^ (0.033 ± 0.013 ng × μL^−1^) for pieces and 0.019 ± 0.005 ng × μL^−1^ from powder. The latter was in the range of the extraction blanks (0.017 ± 0.007 ng × μL^−1^). The quantitation of prepared sequencing libraries by quantitative PCR yielded results in line with the results from DNA extraction (**Figure 2A**).

**Figure 2.**
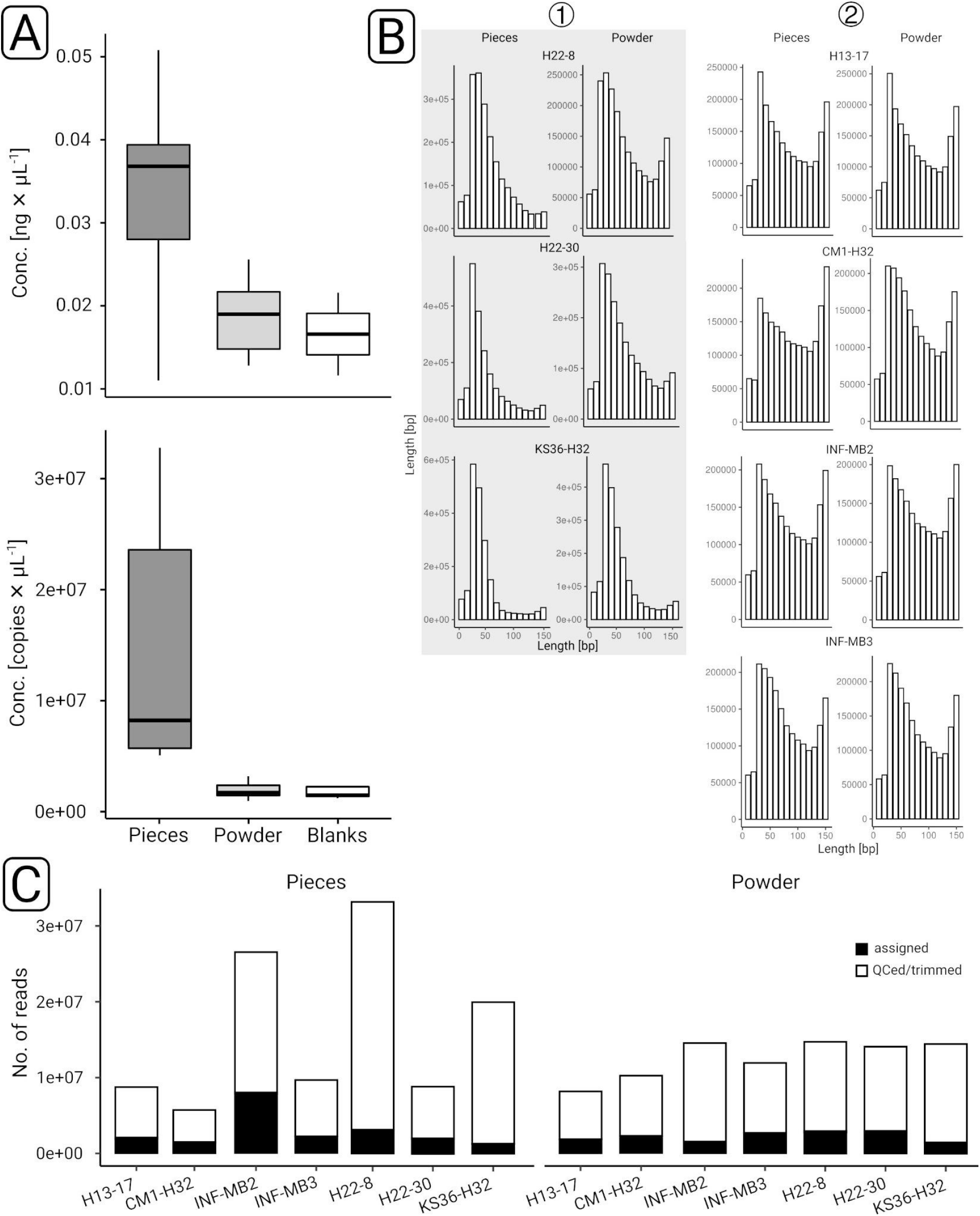
Overview of data (pre-)processing. Samples were quantified by fluorometry and quantitative PCR after DNA extraction (upper panel) and library preparation (lower panel) (A). Sequence length histograms were generated after quality control and trimming based on subsampled (n = 1 M read pairs) data sets. The grey shading highlights three data sets for which the read length distribution was skewed to the left. Based on taxonomic profiling (see main text) we summarized these three data sets in two groups: (1) and (2) (B). The proportion of quality controlled and trimmed sequences that could be assigned taxonomically was determined based on database queries with *diamond* against NCBI nr.

Based on the amount of extracted DNA from the processed samples, we crudely estimated the number of cells potentially present in the rock material. Taking into account the molecular weight of one base pair and using a length of three million base pairs as proxy for a prokaryotic genome, we estimated between 2.81 and 4.25 × 10^5^ cells × g^−1^ processed rock material.

### Taxonomic profiling

Sequence data pre-processing (**Table S1**, **Figure S7**) indicated that the length distribution was generally skewed towards shorter lengths (**Figure 2B**). Consequently, the proportion of taxonomically assigned reads was rather low and varied between 6.2 and 18.6% (**Figure 2C**, **Table S1**). k-mer based redundancy analysis (**Figure S8**) suggested that our data covered more than 90% of the anticipated diversity based on recovered mgDNA. Decontamination analysis identified in total 31 contaminants, one on phylum-level (Spirochaetes), two on class-level (Epsilonproteobacteria, Chlamydiia), 9 on family- and 19 on genus-level (**Table S2**). Principal component analysis on phylum level (**Figure 3A**) showed that H22-8, H22-30, and KS36-H32 were separated from blank data sets, independent from decontamination. The remaining four data sets were grouped together with some of the library and extraction blanks, independent of sample type. Decontamination made data sets more distinguishable from blanks, which was for instance evident in the case of CM1-H32 and H13-17. On family level (**Figure 3B**), decontaminated data sets could be clearly distinguished from blanks. For the subsequent taxonomic profiling pieces and powder data sets have been pooled.

**Figure 3.**
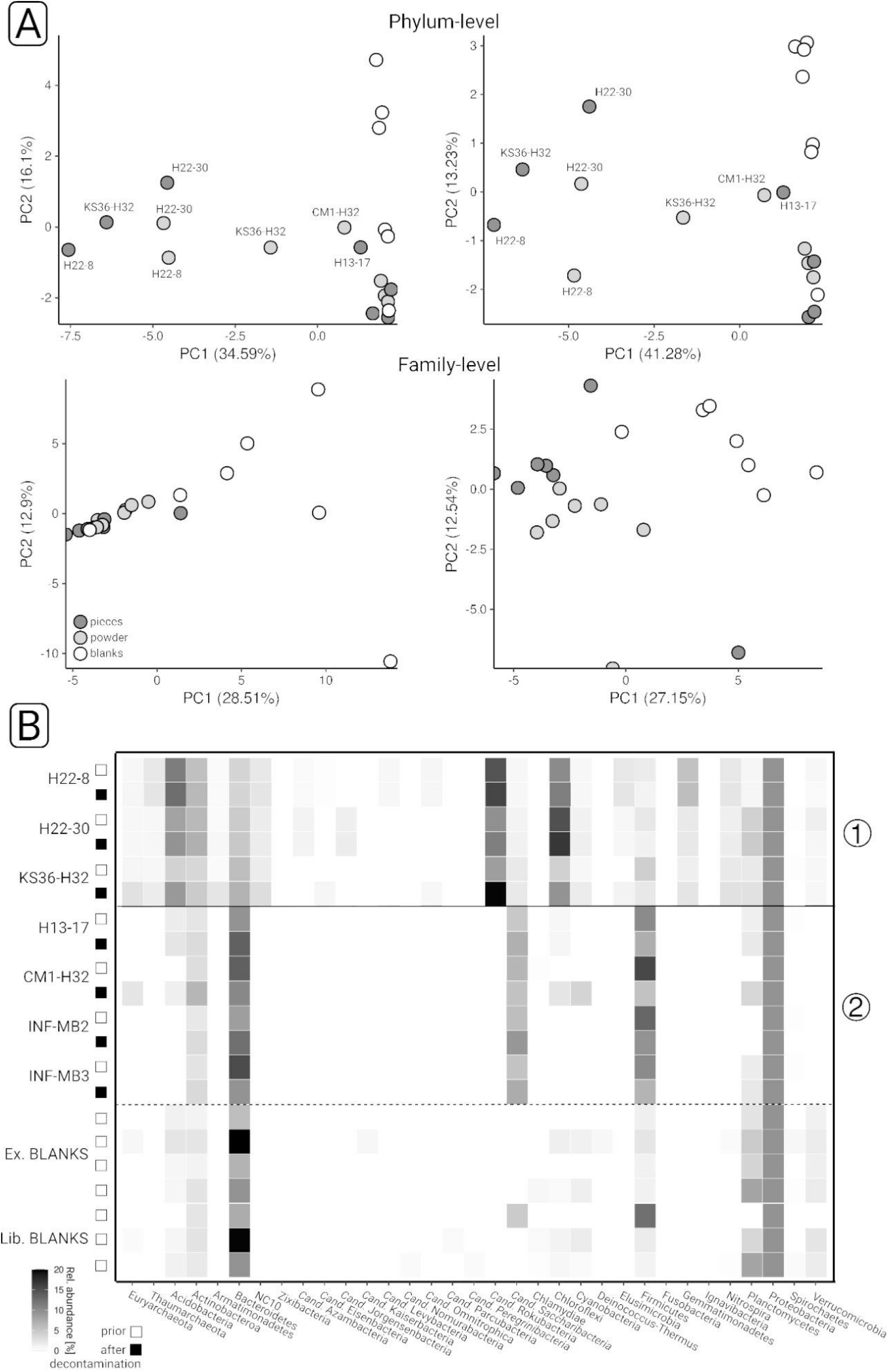
Taxonomic profiling of rock endolithic microbial communities. Principle component analyses were carried out based on phylum-level (A) and family-level (B) taxonomic profiles, prior to (left) and after (right) decontamination. The color coding indicates the sample type. Phylum-level taxonomic profiles were visualized as heatmap (C). (1) and (2) indicate two groups of samples (see main text for details). White and black boxes indicate if the corresponding profile is based on decontaminated data. Ex. and Lib. BLANKS refer to extraction and library blanks, respectively.

Taxonomic profiles were characterized by inverse abundance patterns that divided the data sets into two groups. Group (1) included H22-8, H22-30, and KS36-H32; group (2) H13-17, CM1-H32, INF-MB2, and INF-MB3. Acidobacteria (3.93-11.48%), *Cand.* Rokubacteria (8.28-17.09%), Chloroflexi (4.07-14.74%), Cyanobacteria (0.56-2.71%), NC10 (1.49-4.18%), Nitrospirae (1.33-3.01%), and Thaumarchaeota (0.69-2.75%) (**Figure 3C**, **Table S3**) featured increased abundances in group (1). In comparison, the relative abundances of for instance Firmicutes (up to 15.34%), *Cand.* Saccharibacteria (2.68-9.08%), and Bacteroidetes (15.01-20.08%) were higher in group (2). Some of the mentioned taxa were also detected in the blanks. Bacteroidetes reached abundances up to 36%, while *Cand. Saccharibacteria* were only detected in one blank (4.7%). The relative abundances of Acidobacteria and Chloroflexi did not exceed 2 and 1.5%, respectively. Cyanobacteria abundances were comparable between data sets and blanks. Nitrospirae were only found in two blanks and the abundances were below 0.5% (**Table S3**). Proteobacteria were highly abundant in all data sets (up to 70.81%), and partially much more abundant in the blanks (up to 85.3%).

Decontamination did not lead to major changes in the taxonomic profiles (**Figure 3C**). Lesser abundant phyla increased in relative abundance. Examples include *Cand.* Eisenbacteria (KS36-H32), *Cand.* Jorgensenbacteria (H22-30), *Cand.* Levybacteria (H22-8), and *Cand.* Omnitrophica (KS36-H32, H22-8). Taxonomic profiles at deeper levels are not described as the assignment rate dropped beyond phylum-level.

### Metagenome assembly and DNA damage pattern analysis

For DNA damage pattern analysis, we co-assembled data sets from rock pieces and rock powder from all sites. We compared two different assemblers, *megahit [56]* and *metaSPADES* [58], and ultimately settled on the *megahit* assembly. The assemblies obtained from *metaSPADES* featured larger total assembly lengths, but N50 values and maximum contig lengths were significantly larger when using *megahit* (**Figure S9**). From none of the assemblies, we obtained more than 3153 contigs longer than 1 kbp (1.07-3.15 contigs, 1.81 ± 0.78 [mean ± SD]). The N50 values and the maximum lengths of these contig subsets were rather low, 1.69 ± 0.16 and 16.79 ± 5.43 kbp, respectively. The proportion of recruited reads (after quality control) to the individual assemblies ranged between 6.8 and 23.8% (average 17%), which indicated that our assemblies are only representative for a small part of the generated sequencing data (**Table S4**). We used the mapping files from read recruitment analysis to determine mgDNA fragment lengths (**Figure S10**), which showed that fragment sizes were, with the exception of H22-30, shorter for group (1) samples.

From a taxonomic perspective, the assembled contigs were skewed towards few taxa that assembled well. Contigs from group (2) data sets are dominated by Actinobacteria and Proteobacteria, with combined relative abundances above 95% (**Table S5**). The contigs from group (1) data sets were taxonomically more diverse, but also dominated by Actinobacteria and Proteobacteria, with combined relative abundances of 76% or more. Taxa that were highly abundant based on profiling quality-controlled sequences, were underrepresented. For instance, no more than 0.8% of the assembled contigs were affiliated with *Cand.* Rokubacteria (H22-30) and we only obtained contigs from this taxon from group (1) data sets (**Table S5**).

Mapping metagenomic sequence reads onto assembled contigs larger than 1 kb revealed a pronounced deamination signal in the case of group (1) samples. We detected substitution frequencies partially above 20% (**Figure 4**). Cytosine to thymine substitutions (5pCtoT) and guanine to adenine substitutions (3pGtoA) were comparable for group (1) data sets. Substitution frequencies were negligible for the remaining data sets. The average coverage of the contigs considered for damage analysis was between 65 and 130×, but substantially lower for extraction and library blanks, 24 and 35×, respectively. Extraction and library blanks indicated in comparison to group (1) data sets weak damage signals, with discrepancies between 5pCtoT and 3pGtoA frequencies. The library blanks featured over the first five positions up to 4.2% 3pGtoA, while 5pCtoT did not exceed 1.7% (**Figure 4**). We subsampled contigs affiliated with *Cand. Rokubateria* and detected substitution frequencies between 24 and 32% (**Figure S11**).

**Figure 4.**
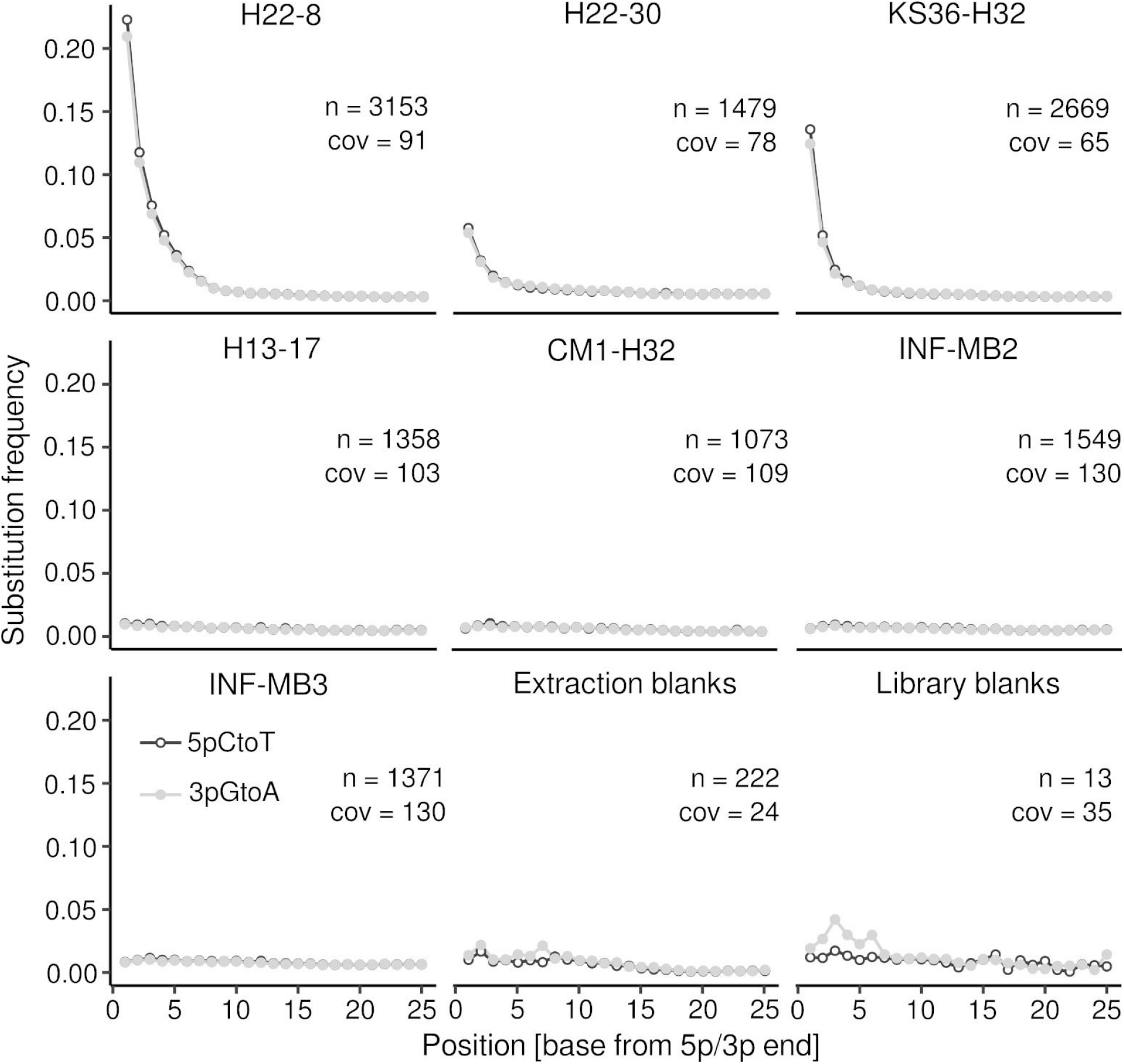
DNA damage pattern analysis. Quality-controlled sequence reads were mapped onto assembled contigs (> 1 kbp). The damage pattern analysis was carried out with *mapdamage* (v.2.2.1) [67]. The plots show the substitution frequency (5pCtoT, 3pGtoA) versus the relative position (from the 5p and 3p end). n = number of contigs > 1kbp considered for the analysis, cov = mean coverage of the contigs.

### Metagenome binning

Metagenome binning led to the reconstruction of 12 bins with a completeness of at least 20% (**Table S6**), five of the reconstructed bins were more than 50% complete. The redundancy of the reconstructed bins was generally low and did not exceed 3.95%, while the heterogeneity reached values of up to 100% (**Table S6**). Nine bins were assigned to *Actinomyces*. Two of the bins belonged to the Acidiferrobacterales (one Sulfurifustaceae [H228_bin5], one Acidiferrobacteraceae [KS36MB2_bin3]). One bin was assigned to UBA9968 (**Table S6**). All of the bins were highly fragmented (no. of contigs > 390), and N50 values did not exceed 4 kbp. In most cases, N50 values were below 2 kbp. The relative abundance of Acidiferrobacterales based on profiling quality-controlled sequences did not exceed 0.28%. They were only detected in H22-8 and KS36-H32. We wanted to compare the two Acidiferrobacterales bins to bins recovered from the Hainich CZE groundwater [72], where this taxon is thought to be involved in sulfur cycling [73], but phylogenomic and ANI (average nucleotide identity)-based comparisons were impossible for the lack of a shared set of single copy marker genes and the high degree of fragmentation.

### Functional profiling

Taking into account that our assemblies recruited only small proportions of the quality-controlled sequences, we used the latter for functional profiling using *humann* [64]. Between 57.3 (INF-MB2) and 85.5% (KS36-H32) (**Figure 5**, “UNMAPPED”) of the sequences did not yield database hits. We regrouped the output from *humann* into KEGG orthologies and summarized the normalized data (copies per million, CoPM) for KEGG pathways (**Table S7**) based on the sequences with database hits. We subsequently focused on pathways that differed between group (1) and group (2) data sets (**Table S8**), in particular functions in the context of carbon fixation, chemolithotrophy, anaerobic respiration, and aromatic hydrocarbon breakdown (**Figure 5**).

**Figure 5.**
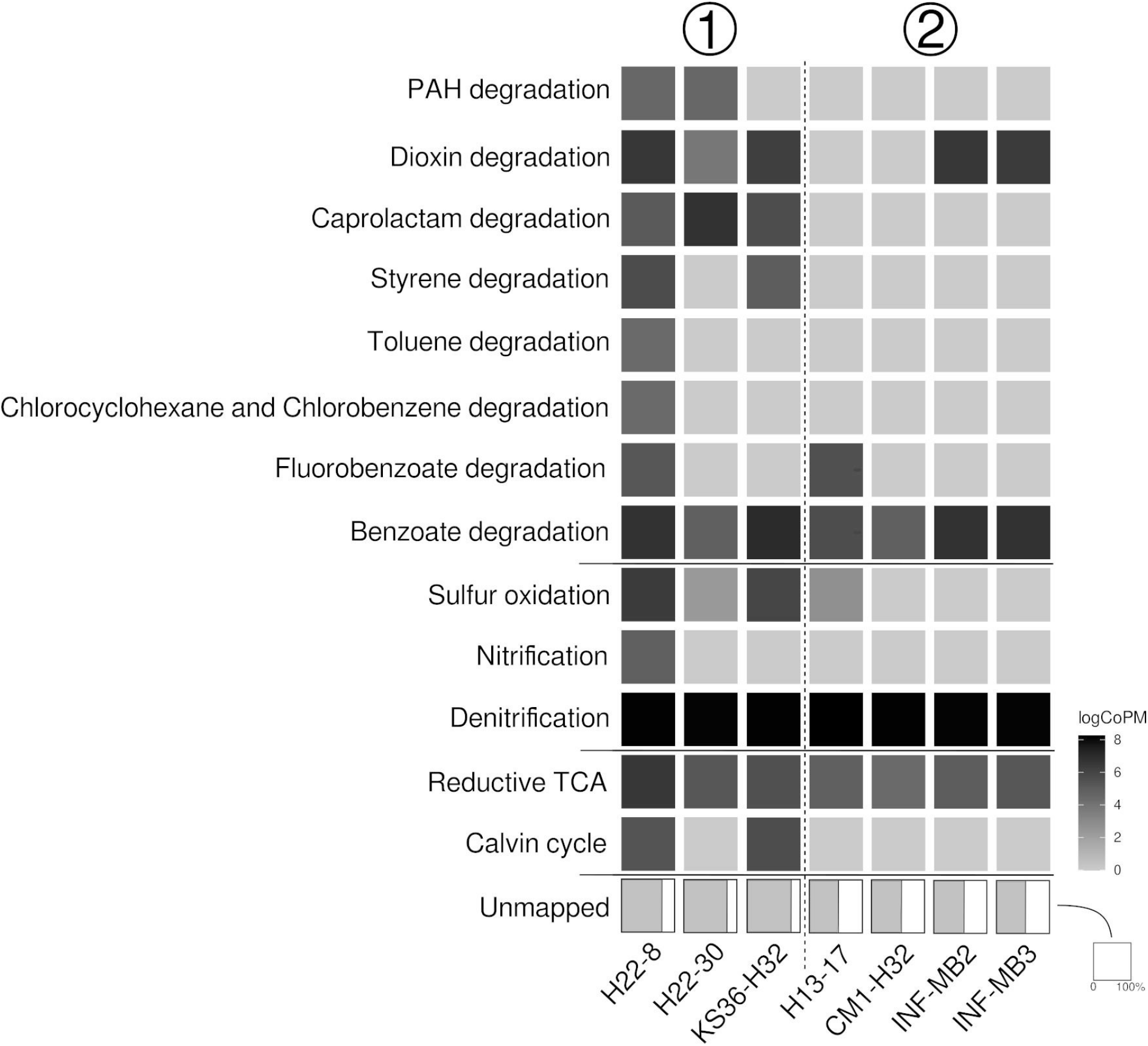
Functional profiling of rock endolithic microbial communities. Profiles were generated based on output from *humann* regrouped into KEGG orthologies. KEGG orthologies were summarized based on pathways and selected functions as described in the methods. Unmapped indicates the proportion of sequences that did not yield any hits against the pre-compiled UniRef databases shipped with *humann*. logCoPM = log copies per million.

Calvin cycle related sequences were only detected for group (1) data sets (H22-8, KS36-H32) that showed pronounced DNA damage. The corresponding logCoPM values were 5.35 and 5.51, respectively. Similarly, evidence for the chemolithotrophic oxidation of sulfur and ammonia was only found in that group, with the exception of H13-17 from group (2). Evidence for nitrification was only found in H22-8. Sequences linked to the reductive TCA cycle were found in all data sets.

Sequences related to aromatic hydrocarbon breakdown were detected in all data sets, with group (1) data sets showing a broader metabolic potential to utilize these substrates, in particular H22-8 (**Table S7 + Table S8**). Matched sequences were affiliated with the breakdown of diverse compound classes, including among others toluene and polycyclic aromatic hydrocarbons (PAH) (**Figure 5**). Group (2) data sets featured comparable narrow metabolic capabilities, including the potential for the breakdown of benzoate and related compounds (**Table S7**).

## Discussion

We were able to recover sufficient mgDNA from all seven rock specimens for metagenomic analysis of endolithic microbial communities using protocols adapted from paleogenomics. The amounts of recovered DNA were extremely small. Extractions from rock pieces were more efficient than from powdered samples. The heat released during powdering may have led to a reduced DNA yield. Estimated cell numbers were within a narrow range of 2.81 and 4.25 × 10^5^ cells × g^−1^ rock, independent from sampling depth and rock characteristics. The subsurface cell count database assembled by Magnabosco and colleagues [3] includes 3787 analyses, of which 2439 were linked to core samples. The database does not include cell counts from limestone, but from rock material classified as carbonate from Lake Van [74] from depths between 0 and 100 mbls (meters below land surface). Our estimated cell numbers lie within the reported broad range (1.27 × 10^3^ - 4.18 × 10^7^ × g^−1^ lake core material). Filling this existing gap is important, given the relevance of carbonate aquifers for global drinking water supply [75]. The majority of the data assembled by Magnabosco and colleagues [3] were based on microscopic counts derived from surface fracture samples after desorbing cells, which might reflect the endolithic community. For obtaining microscopic counts, fluorescent stains like acridine orange or DAPI (4’,6-diamidino-2-phenylindole) are commonly used, which cannot distinguish between dead cells and those with an intact membrane, which are presumably alive. The other fraction of the cell counts was based on qPCR targeting the 16S rRNA gene [3], which is by default also not suited to differentiate between dead and live cells or extracellular DNA. A differentiation between past and potentially alive and active subsurface microbiome members provides relevant information that helps to assess the quality and potential risks associated with groundwater resources. The provision of clean drinking water is considered to be the most important ecosystem service that the subsurface provides to us humans. This service is very vulnerable to anthropogenic and climatic impacts [76].

DNA damage pattern analysis is commonly used in the context of paleogenomics for distinguishing “modern” from degraded “ancient” DNA, which is crucial when studying prehistoric populations of humans, plants, animals, or (pathogenic) microbes [77, 78]. Determined DNA fragment sizes, DNA damage pattern analysis, as well as taxonomic and functional profiling, set apart the group (1) samples. The pronounced damage patterns indicate that DNA obtained from H22-8, H22-30, and KS36-H32 had undergone chemical degradation, which occurs postmortem. The most common forms of DNA damage are depurination, strand breakage, and cytosine deamination on single-stranded overhangs, which occur in sequence during DNA decay [79, 80]. Cytosine deamination occurs at the end of DNA fragments and can be identified by determining the frequency of 5’ cytosine to thymine transitions (3’ guanine to adenine transitions on the reverse complement strand) by mapping metagenomic sequence reads onto metagenome assemblies [66]. We detected substitution frequencies partially above 20%, which is expected for highly degraded DNA from dead organisms [67, 78]. Although, we cannot rule out that all these microbes were already dead when transported into rock matrix pores, it is more likely that they died after being disconnected from energy and water fluxes.

Environmental conditions such as low temperature, high ionic strength, pH, and protection by adsorption can delay the decay of DNA [81–83]. The different forms of crystalline carbonates present in the thin-bedded, alternating mixed carbonate-/siliciclastic bedrock of the Hainich CZE and the INFLUINS site might have favored DNA preservation through neutralizing negative charges, similar to the situation in hard tissue samples (bone, teeth, shells) [28–32]. We propose to consider the genetic records from these three samples as rock paleome signatures, signatures of past microbial communities [84].

Different from sample materials commonly studied for paleogenomics, such as dental calculus, bones, and shells; microbial communities in the subsurface are not necessarily isolated due to being encased by a mineral matrix. Consequenty, microbiome signatures could originate from both, ancient and modern DNA, which affects substitution rates and DNA damage patterns. The subsurface has to be considered as an open system, a giant biogeoreactor with constant or intermittent connection to fluid flow and matter transport, including living microbes [85]. The DNA substitution rates detected for group (1) data sets stress the dominance of decayed DNA in these rock specimens, likely caused by temporary or spatial isolation.

We could not date the DNA due to the tiny amounts recovered. DNA in geological records is in most cases not preserved for more than 10^5^ years [86–90], and 10^6^ years is considered the maximum period over which DNA survival is sufficient for recovery and analysis [91]. The detected paleome signatures cannot reflect the metabolic potentials of microbes colonizing sediments about 240 million years ago, when the Upper Muschelkalk and Lower Keuper (lithostratigraphic subgroups of the Middle Triassic) were formed [92]. Our paleome signatures cannot be considered as biosignatures from ancient microbial life over geological time periods, as those identified in calcite and pyrite veins across the Precambrian Fennoscandian shield by isotopic and molecular analyses [93]. Rather carbonate bedrocks represent DNA archives that can be used to learn more about the near biological past. We argue that distinguishing paleome from non-paleome signatures is a useful approach to identify more recent communities and their functions from those that did contribute to subsurface functioning in the past.

We are confident that the H22-8, H22-30, KS36-H32 data sets are robust. Their taxonomic profiles differed from the laboratory blanks, and they exhibited high DNA fragmentation and higher levels of cytosine deamination than laboratory blanks, indicating that the DNA from group (1) samples disproportionately derives from dead organisms. The remaining “modern” group (2) samples did not feature any pronounced DNA damage and likely originate from alive or recently living organisms.

The paleome signatures of the group (1) samples were all obtained from vadose zone habitats in the low-mountain groundwater recharge area [17]. These shallow bedrock habitats are characterized by spatially and temporally limited water and nutrient supply via seepage from the surface, which can lead to more pronounced starvation especially in disconnected pores compared to saturated habitats. The “modern” signatures of group (2), except H13-17, were obtained from the permanently water-saturated phreatic zone of a fractured aquifer (Hainich CZE) and from ~300 m deep aquitard samples (INFLUINS deep drilling) with similar matrix permeabilities, but without fracture networks [17]. The resulting isolation from the surface did not appear to be critical to the potential survival of endolithic microorganisms in the deep aquitard samples. However, our sample size is too small to conclusively explain the recovery of paleome and non-paleome signatures based on environmental factors or rock characteristics.

Endolithic microbiomes from both groups seem to rely on a bottom-up, chemolithotrophy food web driven by taxa such as *Cand.* Rokubacteria, Gemmatimonadetes, NC10, Nitrospirae, Thaumarchaeota, and Euryarchaeota. Remarkably, we found an increased abundance of chemolithoautotrophs in the paleome signatures coinciding with more detected sequences linked to carbon fixation, nitrification, and sulfur oxidation.

Metagenome assemblies were skewed towards taxa that did assemble well with consequences for DNA damage patterns. Therefore, we also carried DNA damage pattern analysis for only *Cand.* Rokubacteria contigs, and could show that these contigs did feature DNA damage as well, supporting that this taxon was a member of the paleome community. *Cand.* Rokubacteria, was hypothesized to use nitrite oxidation to build a proton motive force [94]. *Cand.* Rokubacteria genomes were previously shown to contain early-branching *dsrAB* genes [95]. They possess motility genes, genes for sensor proteins for diverse stimuli, and genes for respiration (aerobic and anaerobic), fermentation, nitrogen respiration, and nitrite oxidation underline metabolic flexibility and the ability to actively move, which might favor survival in connected rock pore networks.

The phylum NC10, including *Cand.* Methylomirabilis oxyfera, is known to couple anaerobic methane oxidation to nitrite reduction [96]. Nitrospirae and Thaumarchaeota are both well known for nitrification [97, 98], including COMAMMOX in case of the former [99]. Nitrospirae are overall poorly characterized and mostly associated with nitrite oxidation. A candidate genus identified in rice paddy soil, “Candidatus Sulfobium”, was associated with sulfur respiration [100]. Euryarchaeota include the majority of the known methanogens and the Methanosarcinales-related ANME (anaerobic methane oxidizing archaea) clades [101]. However, we cannot make more concrete statements about their specific role in subsurface habitats.

Group (1) data sets showed a broader metabolic potential with respect to sedimentary organic carbon breakdown in the context of aromatic hydrocarbons. The use of sedimentary organic matter by pelagic groundwater microbes of the Hainich CZE was recently shown by DIC isotope pattern analyses [25, 102]. Group (1) samples also featured increased abundances of Acidobacteria. Ubiquitous in soils, Acidobacteria are characterized by a versatility relating to the utilization of (complex) carbohydrates [103] and as K-strategists [104]. Acidobacteria, Bacteroidetes and *Cand.* Saccharibacteria are known as potential degraders for complex polysaccharides [103–106]. The latter two were more abundant in group (2) samples. These taxonomic and metabolic differences suggest a stronger adaptation of the paleome community to the harsh conditions of the endolithic habitat dominated by inorganic electron donors and CO_2_ as carbon source, wheres modern communities might profit from a more constant supply of biomass rich in proteins and carbohydrates under water saturated conditions, which could be derived from plants, but also microbial biomass.

Detected endolithic Cyanobacteria, which have been more prevalent in group (1) samples, could have made use of their fermentative capabilities [107], feeding on available organic carbon, maybe pre-processed by other community members. A study targeting the Iberian Pyrite Belt Mars showed that Cyanobacteria were highly abundant and they seemed to consume hydrogen [15]. Hydrogenotrophy might be a physiological trait in Cyanobacteria dating back to nonphotosynthetic ancestors [108]. Using mgDNA, we detected Candidate Phyla Radiation (CPR) taxa in both groups. We previously hypothesized that CPR taxa are ideally suited to invade and colonize endolithic environments due to their small cell size [17] and their preference to be translocated with seepage water from soil into the vadose zone, and finally into groundwater [109]. This would not apply to episymbiotic CPR with tight relationships with partner organisms. In the paleome, we detected increased abundances of *Cand.* Eisenbacteria, *Cand.* Jorgensenbacteria, and *Cand.* Levybacteria. *Cand.* Eisenbacteria were recently found [110] to possess a potential for secondary metabolite biosynthesis. Because of primer bias of the 16S rRNA gene [111], some CPR may have been missed in many subsurface gene surveys, similar to our previous study of endolithic bacteria from the Hainich CZE [17].

## Summary and conclusion

DNA damage patterns can be used as a proxy to distinguish DNA from intact and potentially alive cells from paleome signatures. Limestone rocks seem to represent ideal archives for genetic records of past microbial communities, due to their specific conditions facilitating long-term DNA preservation. Neither the amount of extractable DNA, nor the status of the endolithic microbiome were indicated by porosity. Water saturation, but not groundwater flow, might be key for microbial survival, as all paleome signatures were detected in the shallow vadose zone, whereas DNA obtained even from deep aquitards, isolated from surface input did not show any DNA decay. Taxonomic and functional profiling highlighted the importance of hydrocarbon utilization and chemolithotrophy linked to sulfur cycling, the latter presumably driven by *Cand.* Rokubacteria in the paleome. Our study shows that carbonate rocks harbor microbial biomass, but that a large portion of the microbes detected by metagenomic sequencing are likely echoes of past microbial communities. Metagenomics and the distinction between “modern” and “ancient” DNA can pave the way to a deeper understanding of the subsurface geomicrobiological history and its changes over time.

## Acknowledgements

This work was supported financially by the Deutsche Forschungsgemeinschaft (DFG, German Research Foundation) under Germany’s Excellence Strategy – EXC 2051 – Project-ID 390713860, the Collaborative Research Centre AquaDiva (CRC 1076 AquaDiva - Project-ID 218627073) of the Friedrich Schiller University Jena, and the Max Planck Society.

## Author contributions

CEW carried out data processing, data analysis, and wrote and revised the manuscript based on input from all co-authors. RS, supported by ZF, was responsible for rock sample processing, testing and adapting protocols, DNA extractions, and sequencing library preparation. IV and AH contributed to sequence data preprocessing, decontamination analysis, and data interpretation. RL coordinated the sampling, acquired permits, and characterized sampled rock material. TR performed μCT analysis. KUT coordinated the sampling, acquired permits, acquired funding, and contributed to data interpretation. CW conceptualized the research, contributed to data interpretation, and acquired funding. KK conceptualized the research, contributed to data interpretation, and acquired funding.

## Supplementary Figures

**Figure S1.**
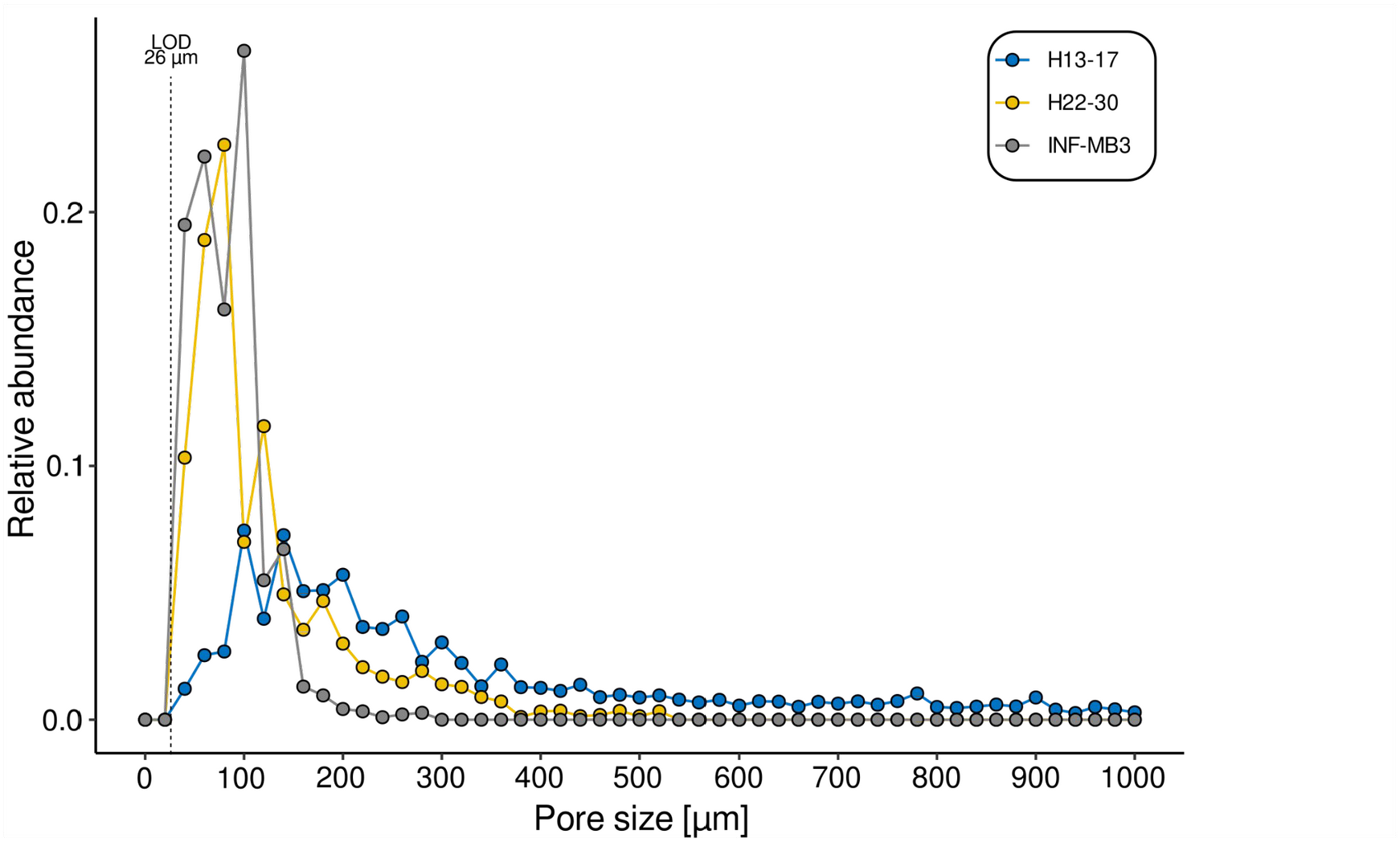

**Figure S2.**
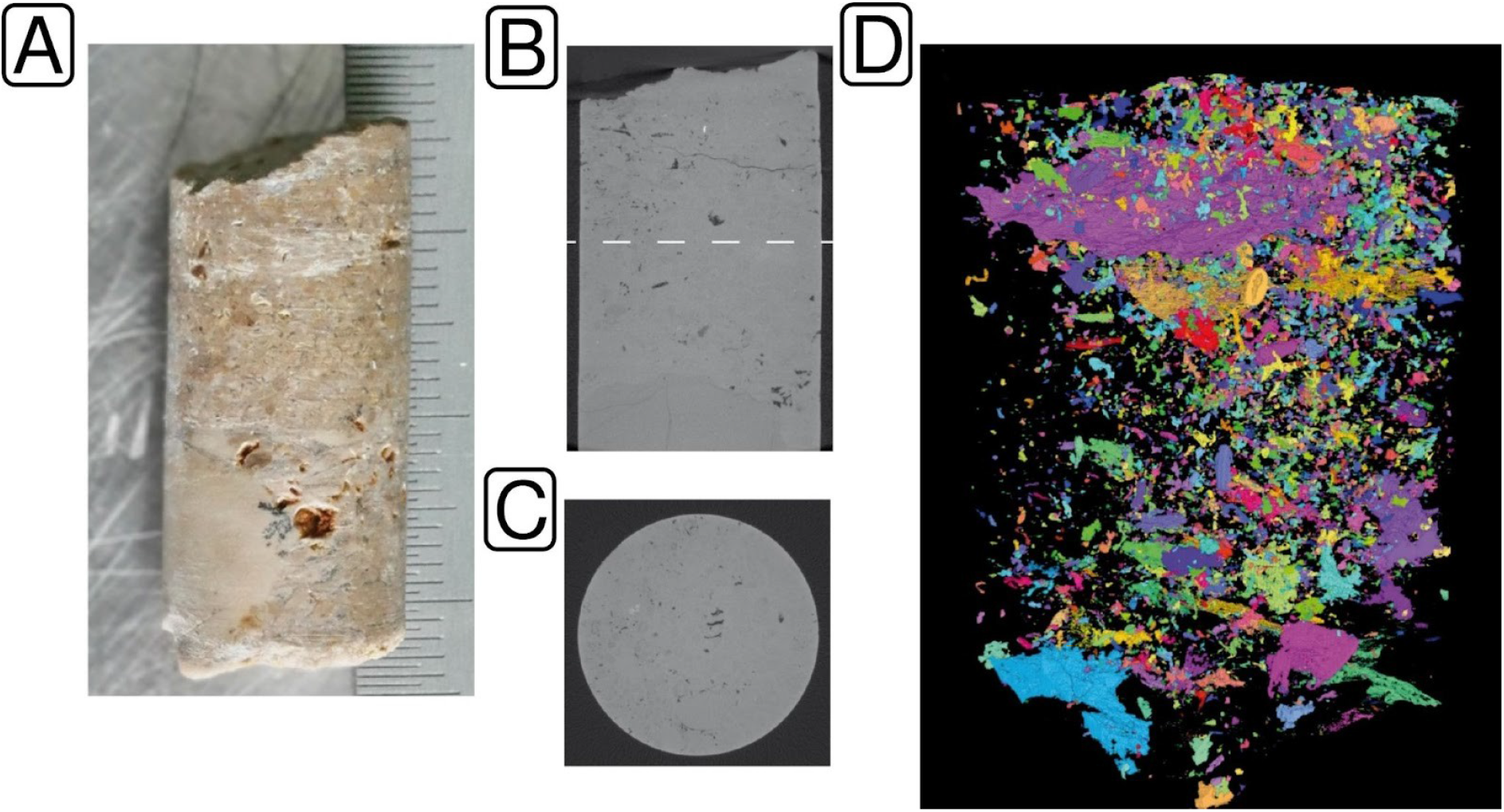

**Figure S3.**
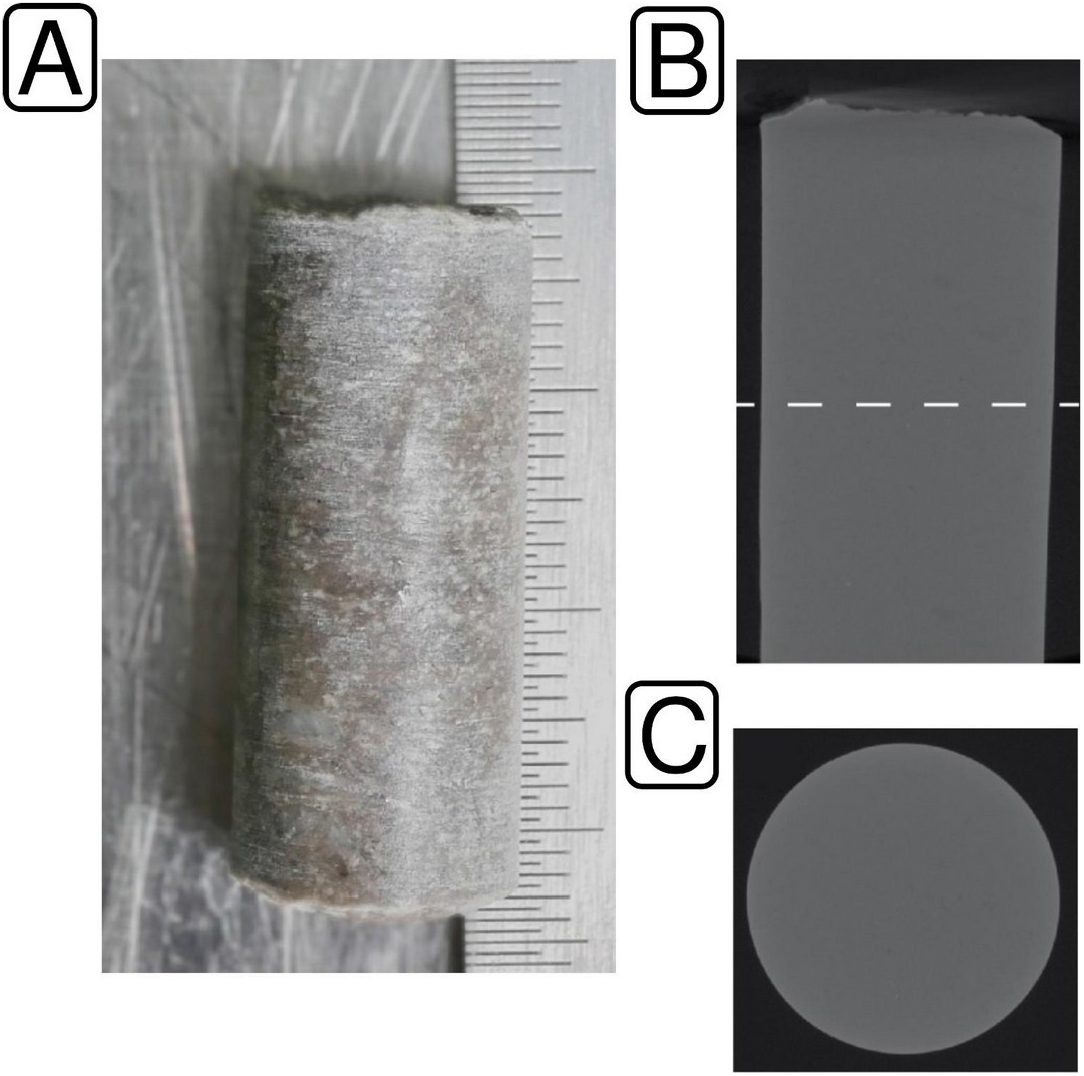

**Figure S4.**
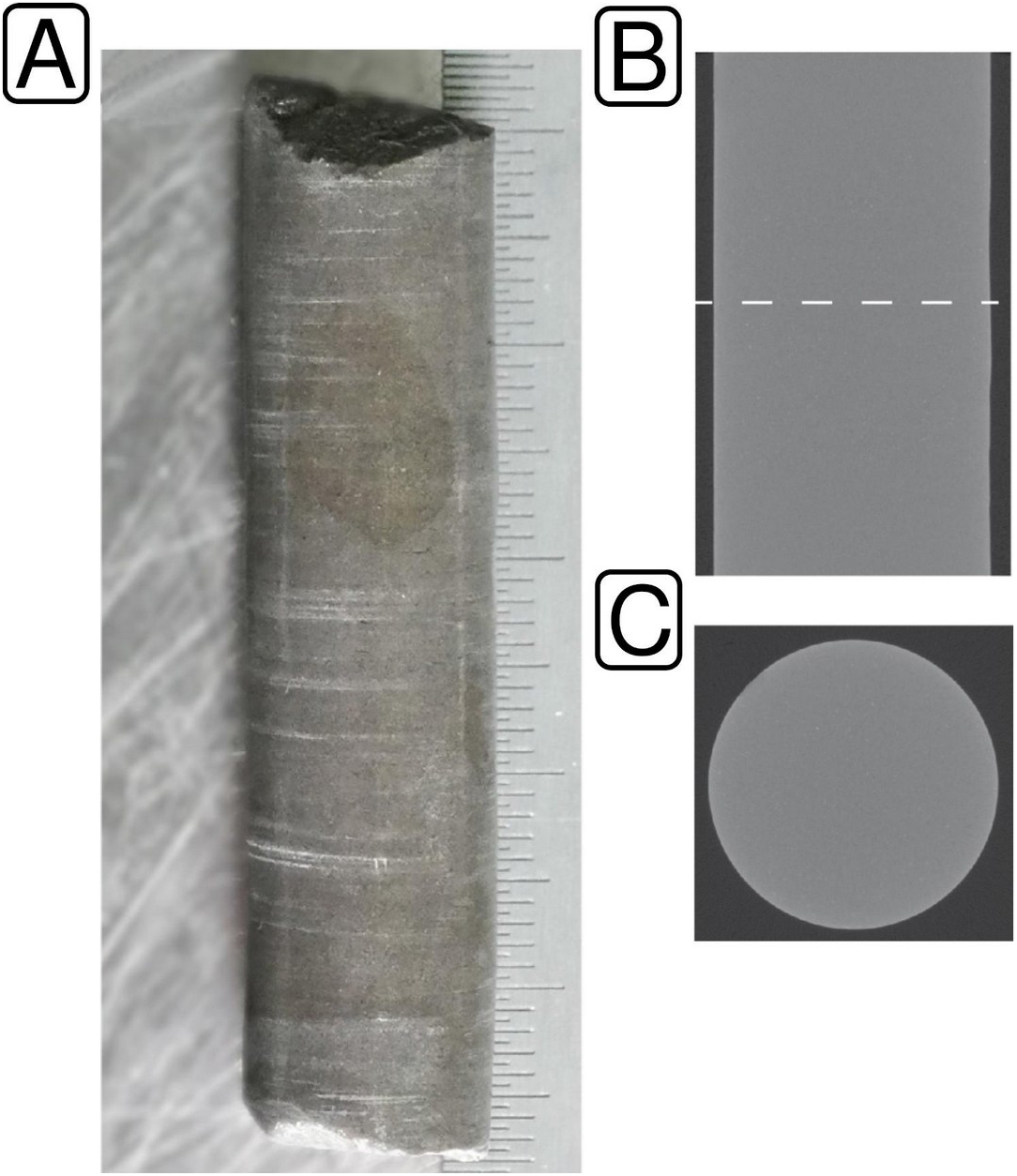

**Figure S5.**
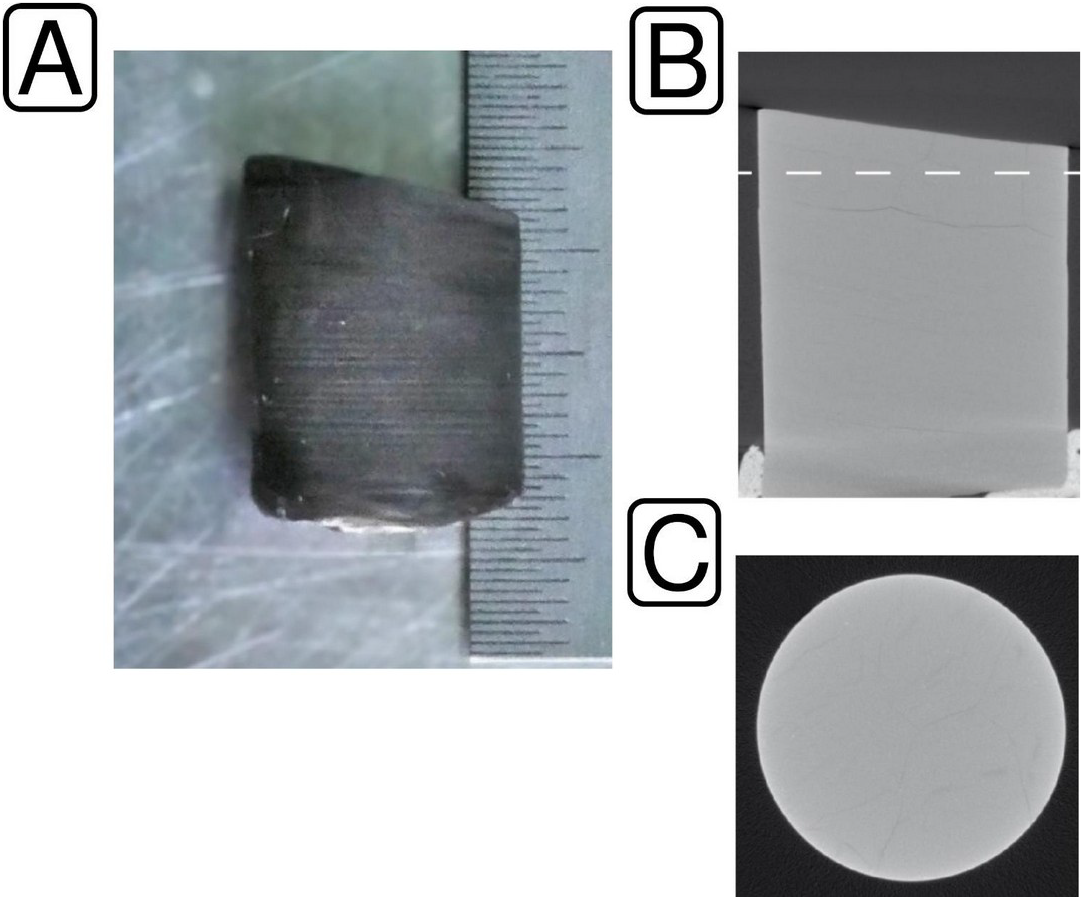

**Figure S6.**
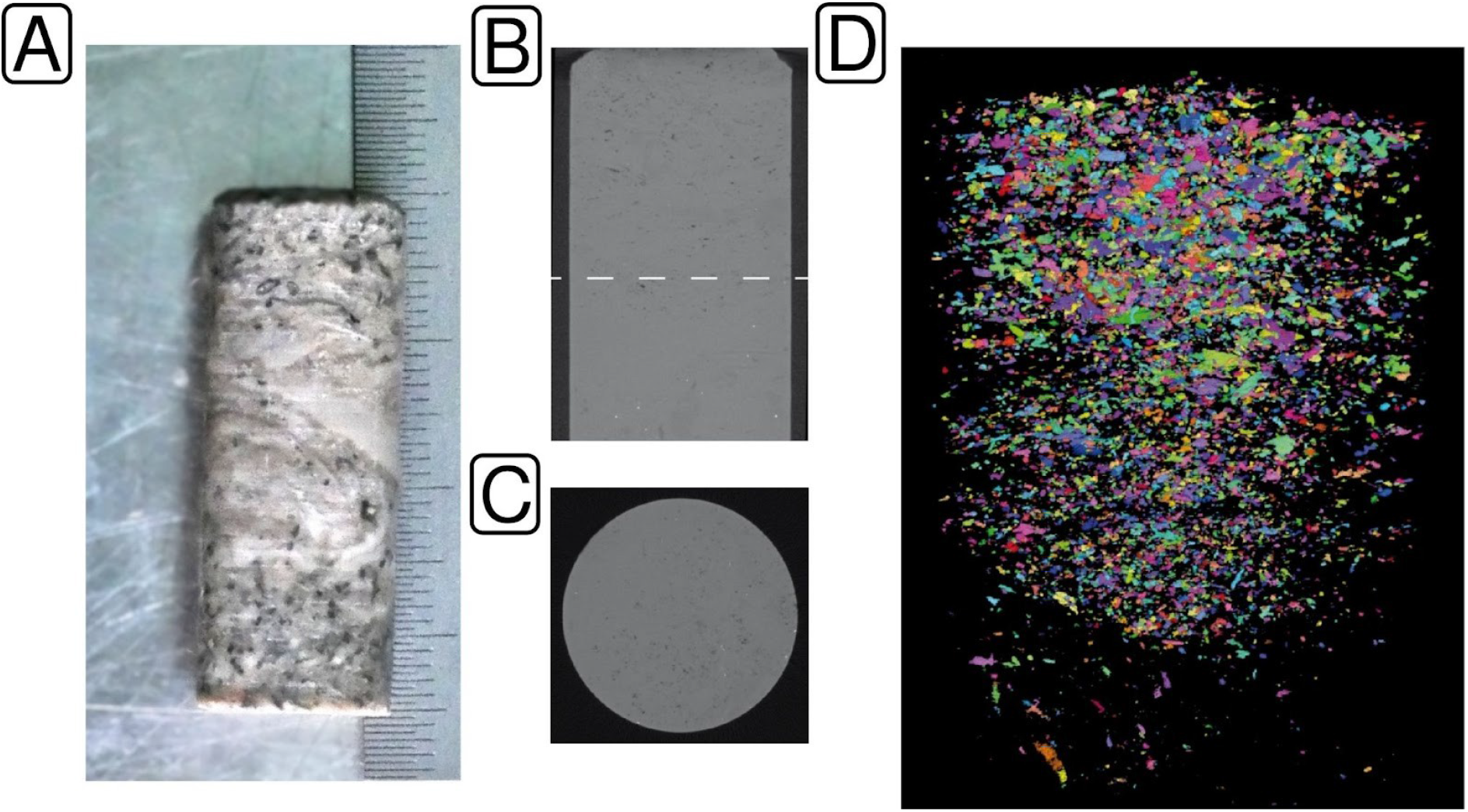

**Figure S7.**
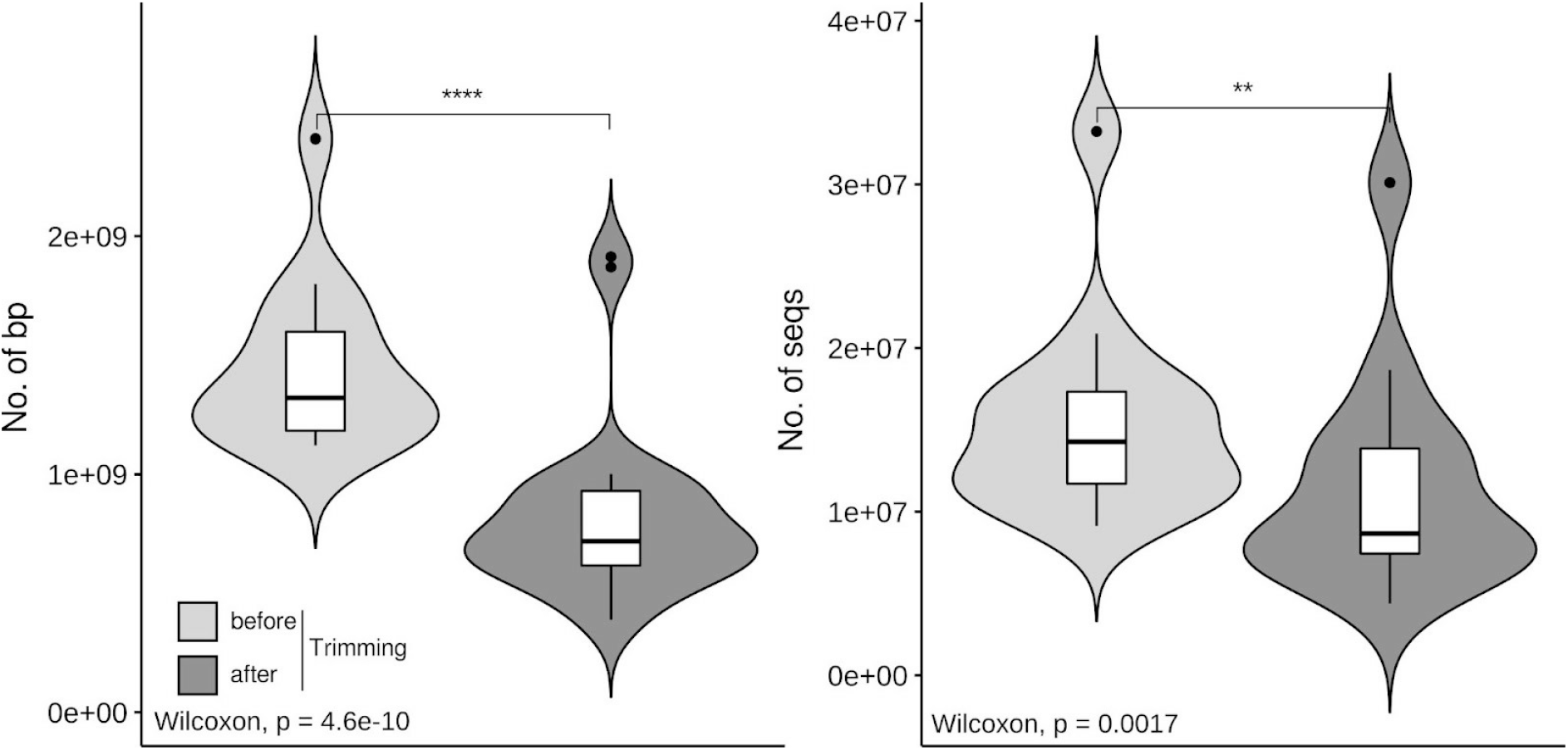

**Figure S8.**
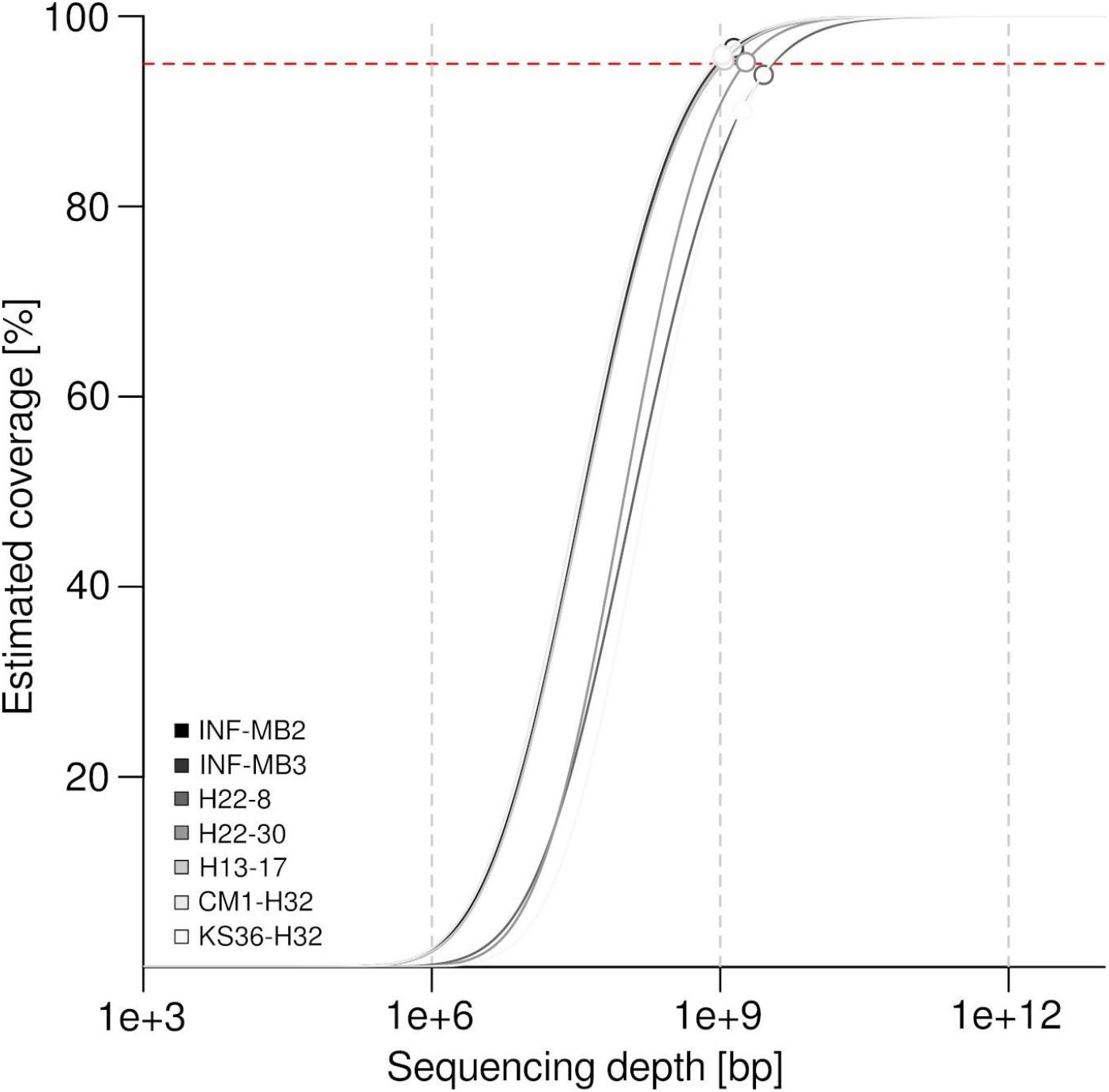

**Figure S9.**
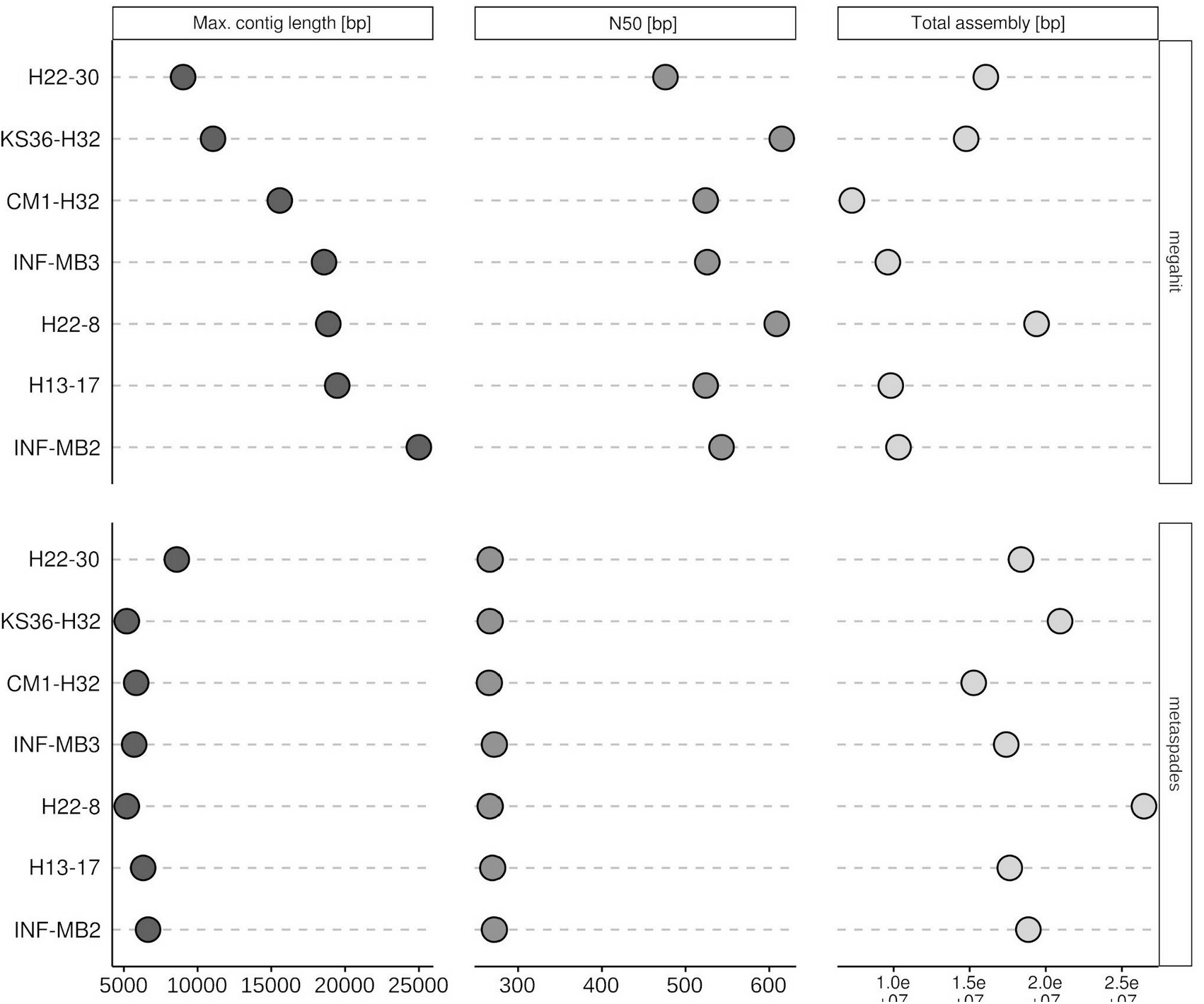

**Figure S10.**
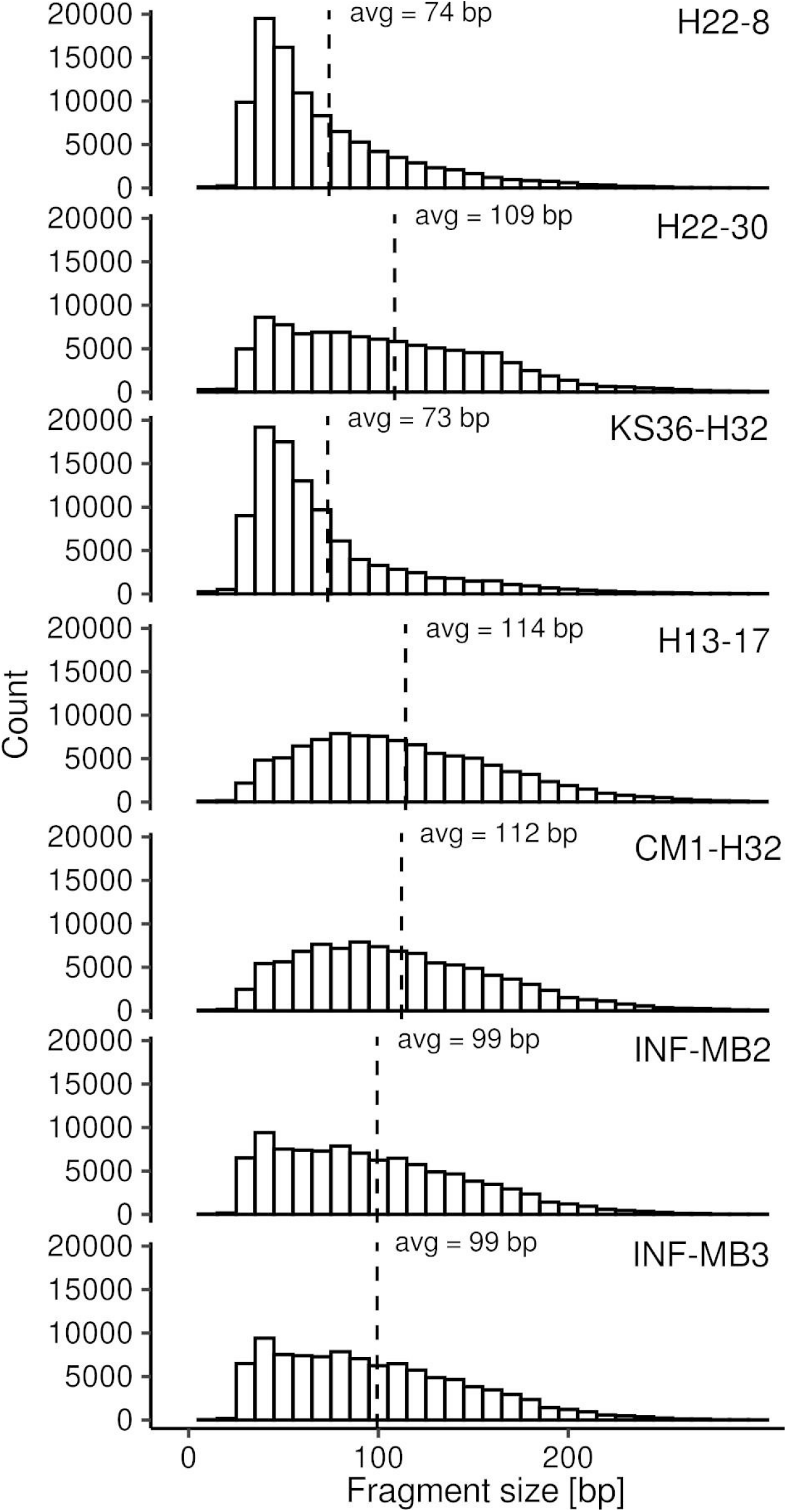

**Figure S11.**
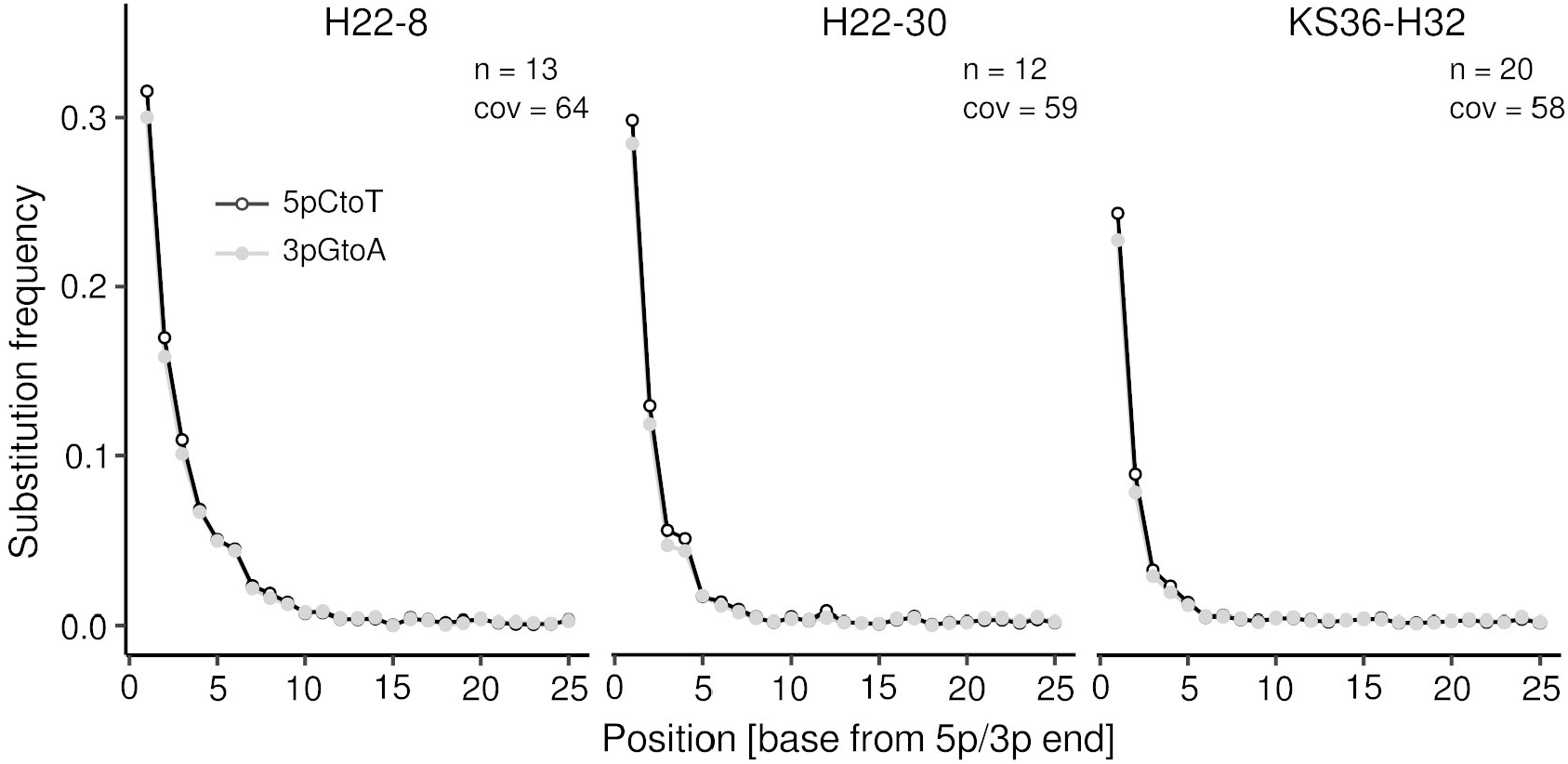

## Notes

### Competing Interest Statement

The authors have declared no competing interest.

