## Supplementary Figures + Table captions for "A glimpse of the paleome in endolithic microbial communities"

### Corresponding author:

Kirsten Küsel

**SUPPLEMENTARY MATERIAL**

**FIG S1.** Pore size distribution of limestone samples with connected pore space (throats >26 µm). LOD = limit of detection.

**FIG S2.** Pore space characteristics of sample H22-30 by µCT analysis. a) Moldic pores (up to large mesopores) dominate over fine fractures in the oolithic packstone. Scale: 0.5 mm. Plug diameter 13 mm. b) Vertical section shows porosity >26 µm. The dashed line marks the position of c. c) Horizontal section shows a moldic pore in a gastropod fossil. d) Reconstructed pore space. Colors mark parts of the pore system connected by throats >26 µm.

**FIG S3.** Pore space characteristics of sample KS36-H32 by µCT analysis. a) The packstone shows minimal alteration (Fe-mineral-stained spots, not visible). Scale: 0.5 mm. Plug diameter 13 mm. b) In the vertical section, connected pores up to ~26 µm (resolution limit) are assumed. The dashed line marks the position of c. c) Horizontal section showing the tight matrix that exhibits no pores connected by throats >26 µm.

**FIG S4.** Pore space characteristics of sample CM1-H32 by µCT analysis. a) The wackestone lacks alteration. Scale: 0.5 mm. Plug diameter 13 mm. b) In the vertical section, connected pores up to ~26 µm (resolution limit) are assumed. The dashed line marks the position of c. c) Horizontal section showing the tight matrix that exhibits no pores connected by throats >26 µm.

**FIG S5.** Pore space characteristics of sample INF-MB2 by µCT analysis. a) The calcareous mudstone lacks alteration. Scale: 0.5 mm. Plug diameter 13 mm. b) In the vertical section, connected pores up to ~26 µm (resolution limit) are assumed. The dashed line marks the position of c. c) Horizontal section showing the tight matrix that exhibits no pores connected by throats >26 µm.

**FIG S6.** Pore space characteristics of sample INF-MB3 by µCT analysis. a) The packstone/grainstone sample lacks alteration. Scale: 0.5 mm. Plug diameter 13 mm. b) Vertical section shows minor porosity >26 µm. The dashed line marks the position of c. c) Reconstructed pore space. Colors mark parts of the pore system connected by throats >26 µm.

**FIG S7.** Number of bp and sequences before and after sequence trimming visualized as a combined box plot and violin plot. Statistical significance was tested using Wilcoxon signed-rank test.

**FIG S8.** Estimated coverage of metagenome data sets based on k-mer based redundancy using *nonpareil* [1,2]. The dashed red line indicates 95% coverage.

**FIG S9.** Assembly statistics for *megahit* [3] and *metaspades* [4] assemblies.

**Fig S10.** DNA fragment size distribution. Fragment sizes were deduced from 100k sampled read pairs mapped onto assembled contigs (> 1 kbp).

**Fig S11.** DNA damage pattern analysis of *Cand.* Rokubacteria contigs. Contigs were subsampled based on the taxonomic affiliation, which was determined with *kaiju* [5]. Quality-controlled sequence reads were mapped onto assembled contigs (> 1 kbp). The damage pattern analysis was carried out with *mapdamage* (v.2.2.1) [6]. "Original" refers to contigs obtained from the *megahit* assembly. Given the atypical damage curves, we tested for problems with consensus sequences from the assembly by correcting the assembly based on read mapping a The plots show the substitution frequency (5pCtoT, 3pGtoA) versus the relative position (from the 5p and 3p end). n = number of contigs > 1kbp considered for the analysis, cov = mean coverage of the contigs.

**Table S1.** Statistics sequence data processing. pwd = powdered sample, pc = rock pieces sample, QC = quality control.

**Table S2.** Identified contaminants for different taxonomic ranks. Freq = frequency, prev = prevalence, p.freq = tail probability at value R, p.prev = tail probability of the chi-square distribution for the respective taxon based on presence/absence in true samples and negative controls, p = p-value from Fisher's exact test, NA = not available. Please see [7] for details regarding the mentioned metrics.

**Table S3.** Phylum-level taxonomic profiles. lib_blk = library blank, ex_blank = extraction blank, pc = rock pieces sample, pwd = powdered sample.

**Table S4.** Basic assembly statistics and results from read recruitment.

**Table S5.** Phylum-level taxonomic profiles of assembled contigs (> 1 kbp).

**Table S6.** Taxonomy and quality information regarding recovered genome bins based on *checkm* [8] output.

**Table S7:** Functional profile based on KEGG Lvl3 Orthologies. Abundances are given as CoPM. CoPM = copies per million.

**Table S8:** Functional profile based on a subset of KEGG pathways. CoPM = copies per million, logCoPM = log copies per million.

**Note:** All supplementary tables are provided in one combined spreadsheet.

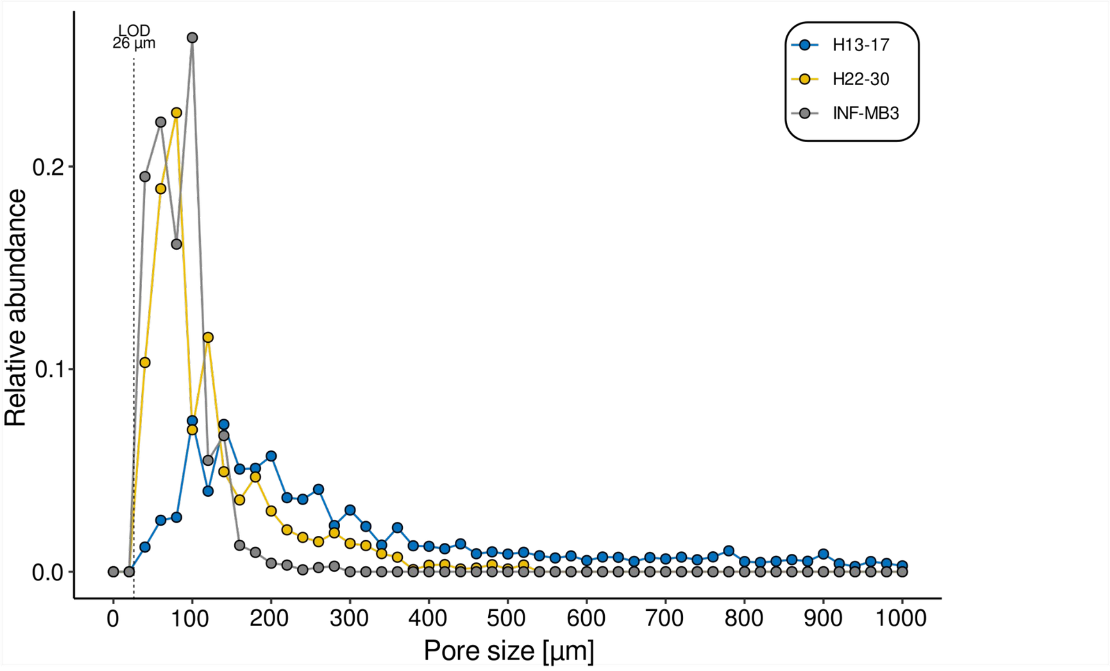

LOD
26 µm
H13-17
H22-30
INF-MB3
Relative abundance
0.0
0.1
0.2
0
100
200
300
400
500
600
700
800
900
1000
Pore size [µm]


**Figure S1**

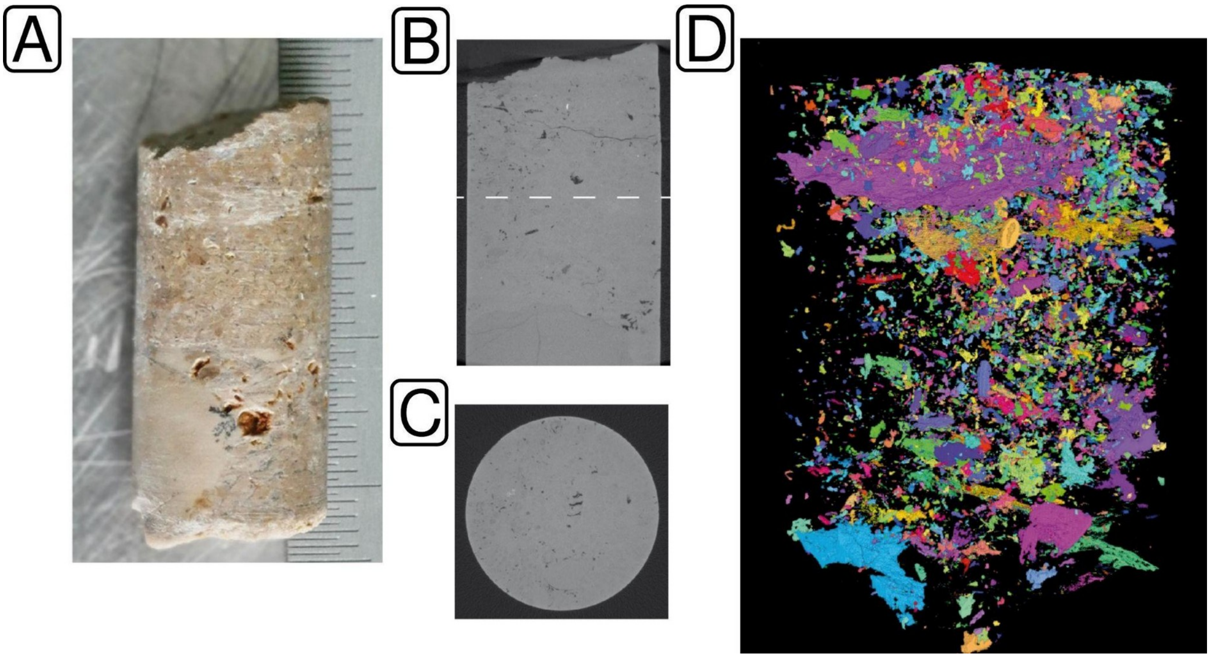


**Figure S2**

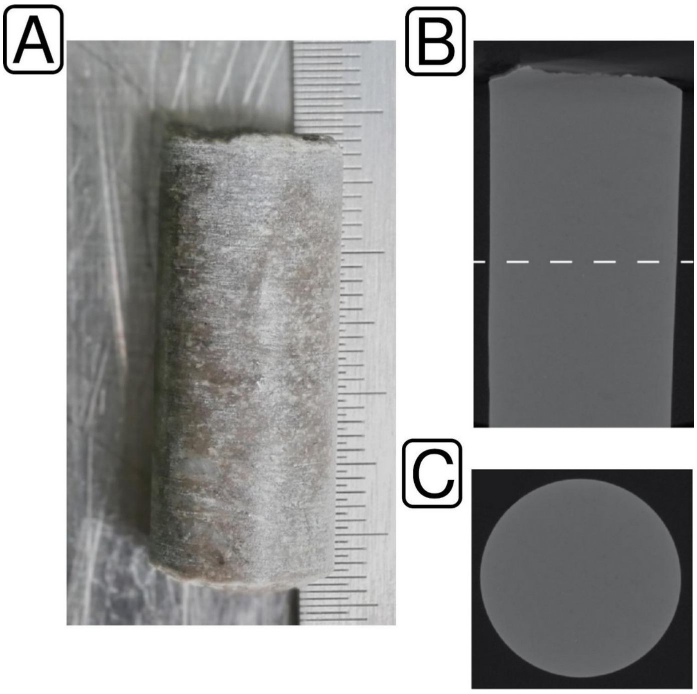


**Figure S3**

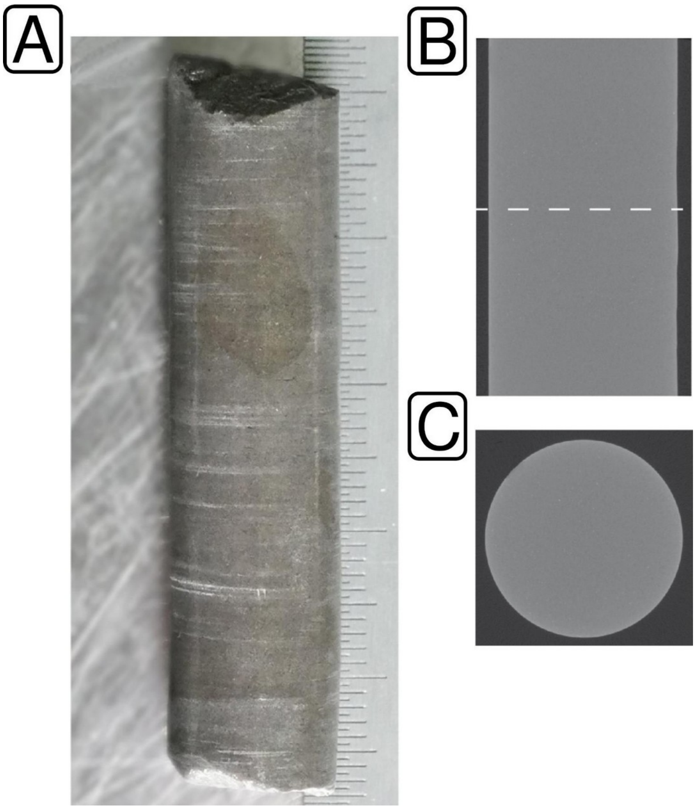


**Figure S4**

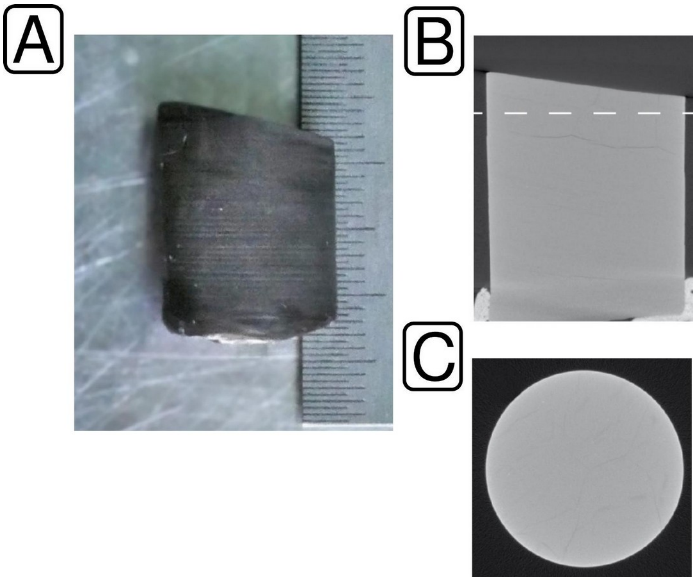


**Figure S5**

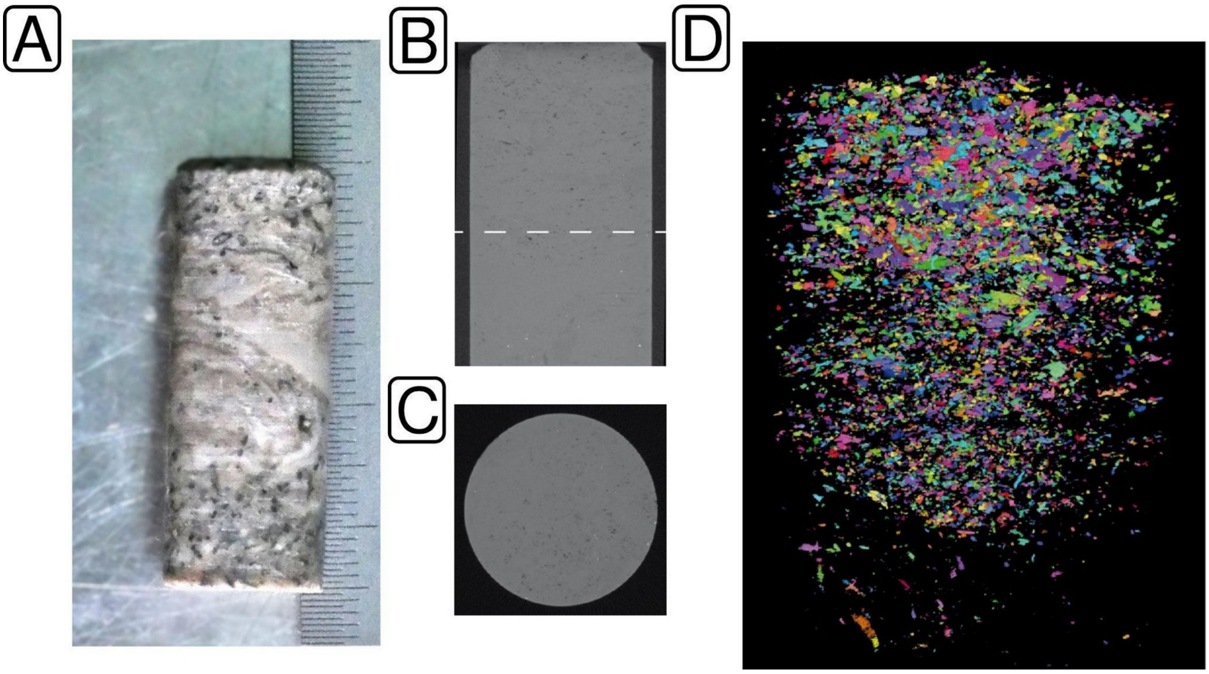


**Figure S6**

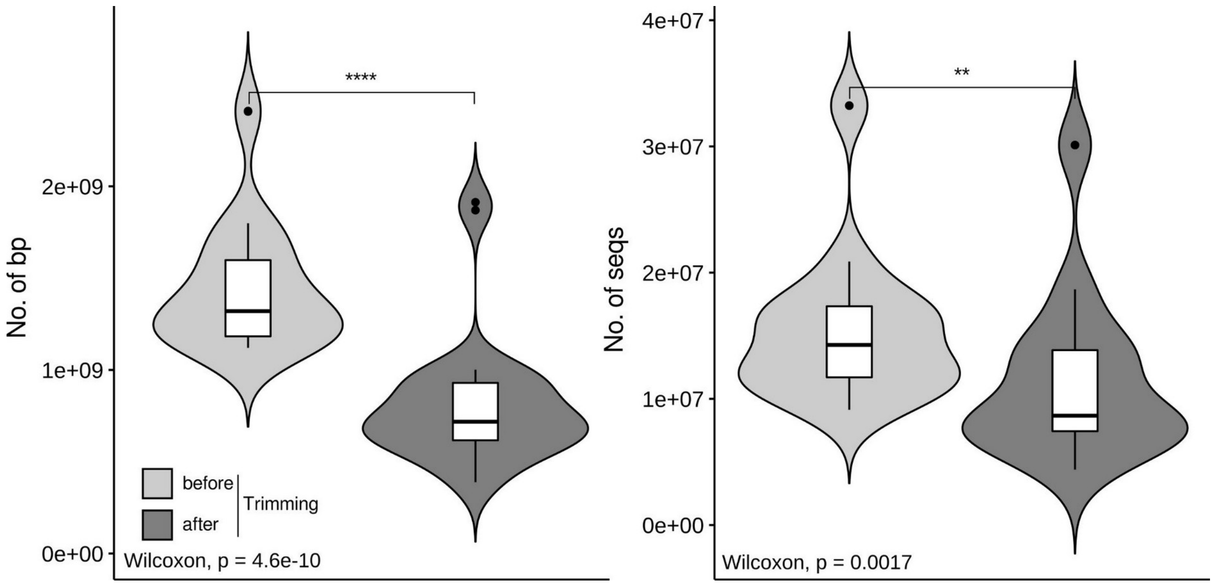

No. of bp
2e+09
1e+09
0e+00
****
before
after
Trimming
Wilcoxon, p = 4.6e-10
No. of seqs
4e+07
3e+07
2e+07
1e+07
0e+00
**
Wilcoxon, p = 0.0017


**Figure S7**

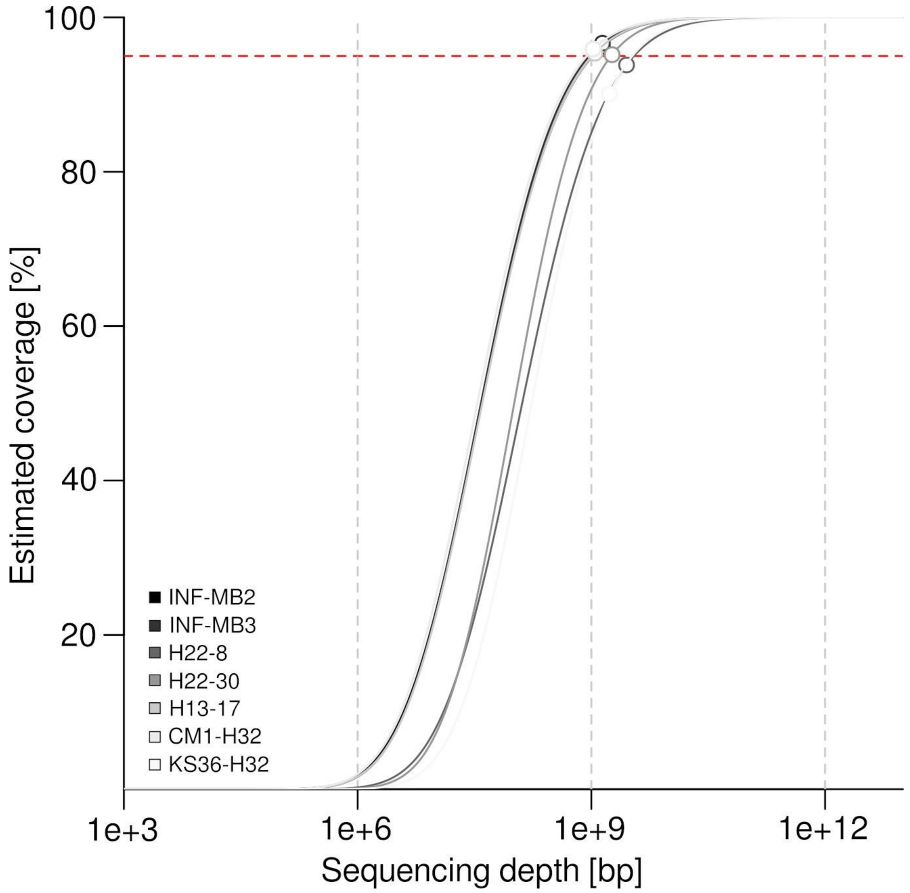

100
80
60
40
20
Estimated coverage [%]
INF-MB2
INF-MB3
H22-8
H22-30
H13-17
CM1-H32
KS36-H32
1e+3
1e+6
1e+9
1e+12
Sequencing depth [bp]


**Figure S8**

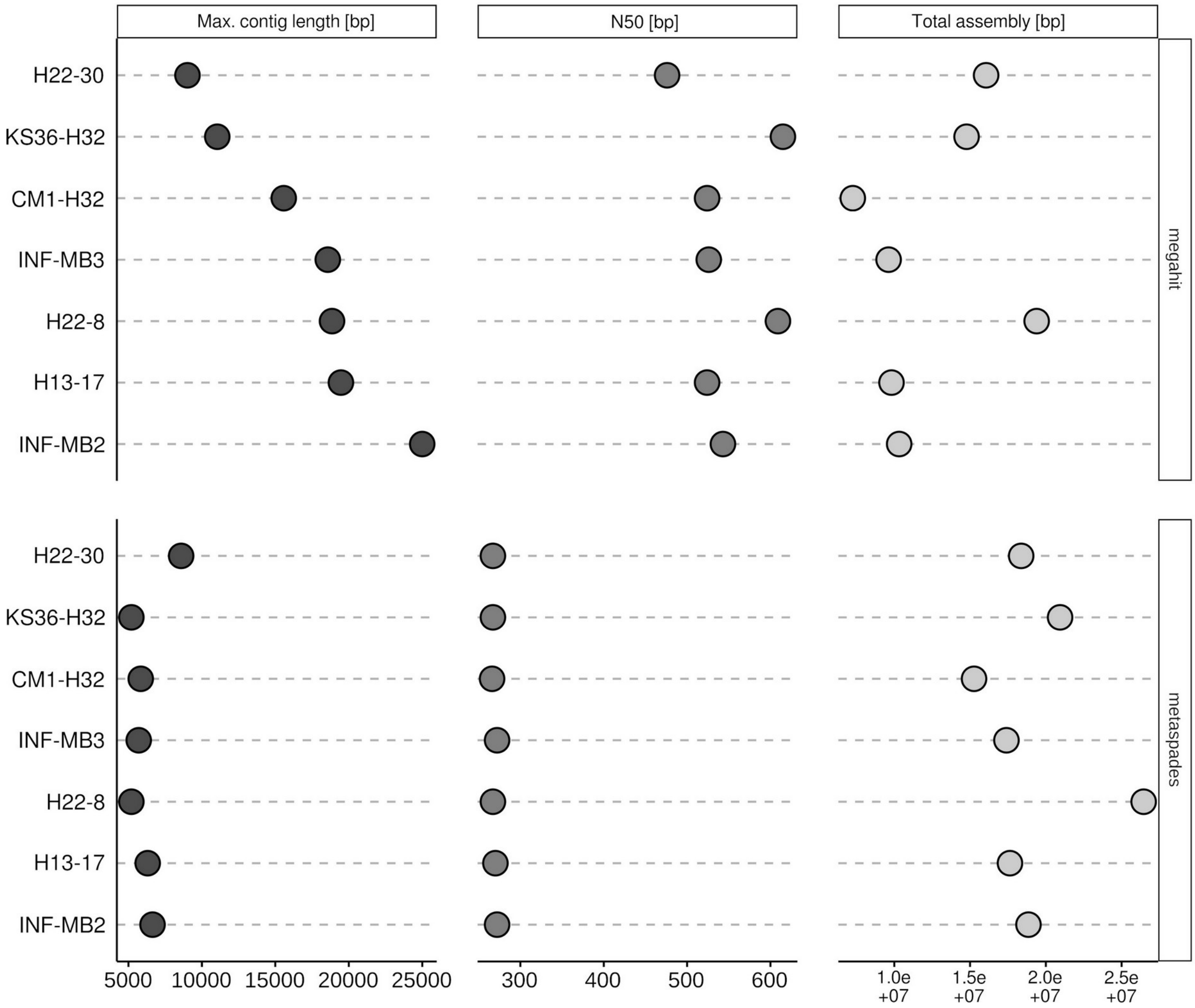
Max. contig length [bp]
N50 [bp]
Total assembly [bp]
megahit
metaspades
H22-30
KS36-H32
CM1-H32
INF-MB3
H22-8
H13-17
INF-MB2
5000 10000 15000 20000 25000
300 400 500 600
1.0e+07 1.5e+07 2.0e+07 2.5e+07

**Figure S9**

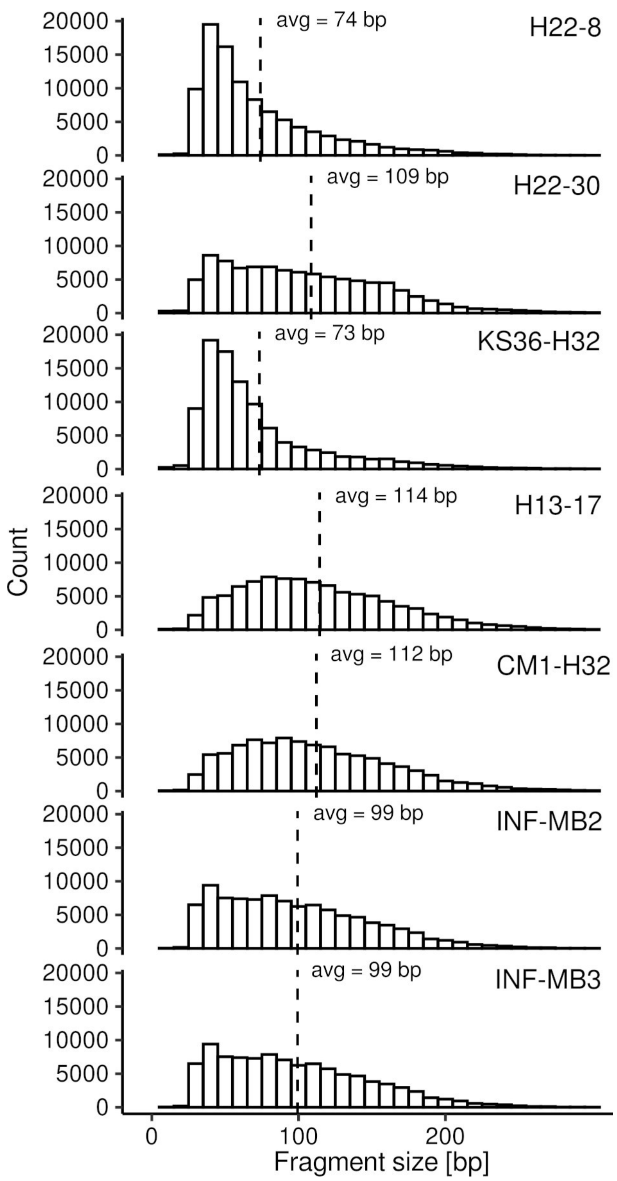

Count
Fragment size [bp]
0
100
200
20000
15000
10000
5000
0
avg = 74 bp
H22-8
avg = 109 bp
H22-30
avg = 73 bp
KS36-H32
avg = 114 bp
H13-17
avg = 112 bp
CM1-H32
avg = 99 bp
INF-MB2
avg = 99 bp
INF-MB3


**Figure S10**

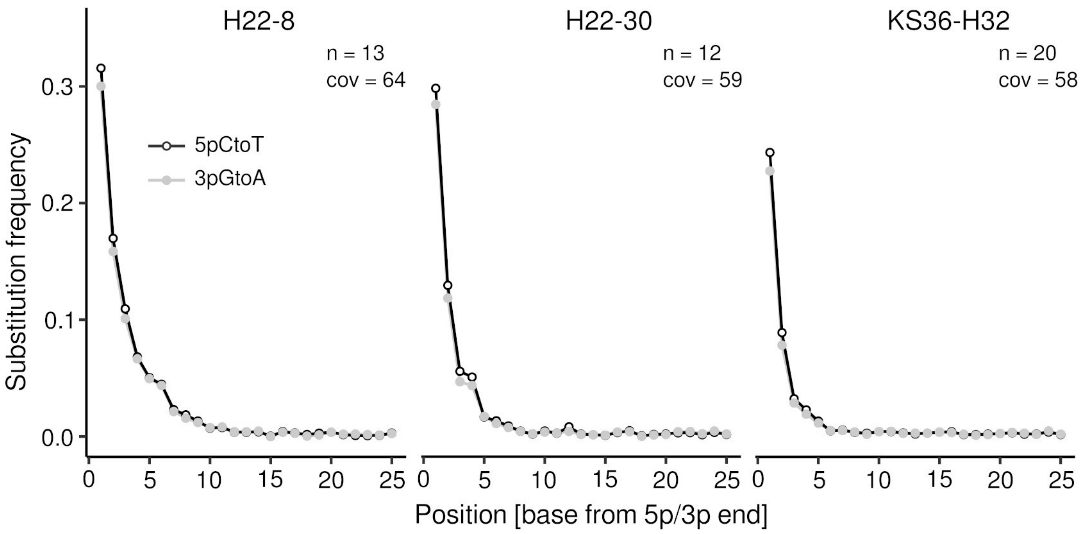
H22-8
H22-30
KS36-H32
n = 13
cov = 64
n = 12
cov = 59
n = 20
cov = 58
5pCtoT
3pGtoA
Substitution frequency
0.3
0.2
0.1
0.0
0
5
10
15
20
25
Position [base from 5p/3p end]

**Figure S11**
